# Dorsal and ventral mossy cells differ in their long-range axonal projections throughout the dentate gyrus of the mouse hippocampus

**DOI:** 10.1101/2020.09.27.315416

**Authors:** Justin J. Botterill, Kathleen J. Gerencer, K. Yaragudri Vinod, David Alcantara-Gonzalez, Helen E. Scharfman

**Author notes:** **Submitting and Corresponding Author:** Helen E. Scharfman, Center for Dementia Research, The Nathan Kline Institute, 140 Old Orangeburg Rd. Bldg. 35, Orangeburg, NY, 10962.

## Abstract

Glutamatergic hilar mossy cells (MCs) have axons that terminate both near and far from their cell body but stay within the DG, making synapses in the inner molecular layer primarily. The long-range axons are considered the primary projection, and extend throughout the DG ipsilateral to the soma, and project to the contralateral DG. The specificity of long-range MC axons for the inner molecular layer (IML) has been considered to be a key characteristic of the DG. In the present study we made the surprising finding that dorsal MC axons are an exception to this rule. We used two mouse lines that allow for Cre-dependent viral labeling of MCs and their axons: dopamine receptor d2 (Drd2-Cre) and calcitonin receptor-like receptor (Crlr-Cre). A single viral injection into the dorsal DG to label dorsal MCs resulted in labeling of MC axons in both the IML and middle molecular layer (MML). Interestingly, this broad termination of MC axons applied to all long-range axons. In contrast, long-range axons of ventral MCs mainly terminated in the IML, consistent with the literature. Taken together, these results suggest that dorsal and ventral MCs differ significantly in their axonal projections, and the difference is primarily in their long-range projections. Since those projections are thought to terminate primarily on GCs, the results suggest a dorsal-ventral difference in MC activation of GCs. The surprising difference in dorsal and ventral MC projections should therefore be considered when evaluating dorsal-ventral differences in DG function.

## 1. INTRODUCTION

The dentate gyrus (DG) of the hippocampus is considered critical in cognitive and behavioral functions. It also has been implicated in several neurological and psychiatric conditions (Scharfman, 2007b). Dentate granule cells (GCs) are the main excitatory cell type, and form a key relay from the entorhinal cortex to area CA3 (Amaral et al., 2007). Inhibitory GABAergic interneurons in the DG provide the main source of inhibition to GCs (Houser, 2007). Hilar mossy cells (MCs) are large glutamatergic neurons that innervate both GCs and inhibitory GABAergic neurons within the DG (Scharfman, 2016; Scharfman and Myers, 2012). MCs make up the majority of hilar neurons, and are known for their complex spines called thorny excrescences (Scharfman, 2016; Scharfman and Myers, 2012). They have dendrites mainly in the hilus and their axon projects to locations within the DG. Near the cell body the axon makes collaterals that terminate mainly in the hilus. Distal to the cell body the axon terminates at many septotemporal levels. There is also a commissural projection that terminates in the contralateral DG (Scharfman and Myers, 2012). The complex projections of MCs have led to considerable interest in their contribution to DG function.

Numerous studies have documented the MC axon projection (Scharfman and Myers, 2012), but a seminal study used biocytin to label individual MCs *in vivo* and quantify the axon projections (Buckmaster et al., 1996). That study found that while ∼25% of the MC axon is located in the hilus, over 60% of the axon was located in the molecular layer (ML). Consistent with prior studies, the majority of the MC axon projected to the inner molecular layer (IML). However, a small fraction of the axon was found in the middle molecular layer (MML) and minimal expression was found in the outer molecular layer (OML). Using electron microscopy, the authors showed that the primary target of long-range MC axons are GCs, supporting the view that MCs primarily activate GCs (Buckmaster and Schwartzkroin, 1994; Buckmaster et al., 1996).

Historically, MCs have been challenging to study due to the lack of specific tools to visualize or manipulate their activity (Scharfman, 2017; Scharfman and Myers, 2012). Technical advances over the past several years have generated specific transgenic mouse lines and viral methods to label MCs and their axons with a high degree of specificity (Scharfman, 2016, 2017). Two of the most widely used mouse lines to study MCs include calcitonin receptor-like receptor (Crlr-Cre) and dopamine receptor d2 (Drd2-Cre) mice. The robust nature of Cre-dependent viral labeling in both of these lines is well documented (Azevedo et al., 2019; Botterill et al., 2019; Jung et al., 2019; Oh et al., 2019; Puighermanal et al., 2015; Senzai and Buzsaki, 2017; Yeh et al., 2018). Several studies have now used these Cre lines to evaluate effects of MC manipulations *in vivo* or *in vitro* (Azevedo et al., 2019; Bernstein et al., 2020; Botterill et al., 2019; Jinde et al., 2012; Jung et al., 2019; Oh et al., 2019; Puighermanal et al., 2015; Senzai and Buzsaki, 2017; Yeh et al., 2018). However, these mouse lines are also useful to address the MC axon projections in the adult mouse. This type of investigation is valuable because past studies mainly used rats, and in addition there is excellent identification of membrane processes using viral labeling of proteins that insert into plasma membranes (Lanciego and Wouterlood, 2020).

In the present study we utilized Cre-dependent viral labeling to evaluate the long-range axons of MCs in Crlr-Cre and Drd2-Cre mouse lines. We administered a single viral injection into the dorsal or ventral hilus to determine whether dorsal and ventral MCs differ in their long-range projections. Both dorsal and ventral injections labeled a large number of MCs proximal to the injection site as well as long-range MC axons throughout the septotemporal axis of the DG bilaterally. Surprisingly, dorsal MCs had a remarkably different pattern of long-range axonal projections compared with ventral MCs. Specifically, the axons that targeted distal locations terminated in both the IML and MML, with a small degree of labeling in the OML. This pattern occurred in both distal ipsilateral and contralateral DG. In contrast, a single ventral injection of virus labeled ventral MCs with axons that were primarily restricted to the IML, consistent with past studies. Taken together, this study provides novel evidence that dorsal and ventral MCs differ in their anatomical projections and these findings should be considered when evaluating how MCs influence the activity of the DG network.

## 2. METHODS

### 2.1 Animals and genotyping

All experimental procedures were done in accordance with the National Institutes of Health (NIH) guidelines and approved by the Institutional Animal Care and Use Committee (IACUC) at the Nathan Kline Institute. Adult male and female Drd2-Cre^+/-^ and Crlr-Cre^+/-^ mice were used in the present study (age range: 2-5 months). Hemizygous Drd2-Cre and Crlr-Cre males were bred in-house to wild-type C57BL/6 females. Breeding pairs were fed Purina 5008 rodent chow (W.F. Fisher) and provided 2”x2” nestlets (W.F. Fisher). Mice were weaned at postnatal day 25-30 and housed with same-sex siblings (2-4 per cage) in standard laboratory cages with corn cob bedding. Mice were maintained on a 12 hr light-dark cycle with standard rodent chow (Purina 5001, W.F. Fisher) and water available *ad libitum*. Genotyping was performed by the Genotyping Core Laboratory at New York University Langone Medical Center.

### 2.2 Stereotaxic surgery and viral injections

Stereotaxic surgery was performed as previously described (Botterill et al., 2019). Mice were anesthetized with isoflurane (5% induction, 1-2 % maintenance; Aerrane, Henry Schein) and secured in a rodent stereotaxic apparatus (Model #502063, World Precision Instruments). Buprenex (Buprenorphine, 0.1 mg/kg, s.c.) was delivered prior to surgical procedures to reduce discomfort. Body temperature was maintained at 37 °C via a homeothermic blanket system (Harvard Apparatus). The scalp of each mouse was shaved and swabbed with betadine (Purdue Products) and lubricating gel was applied to the eyes to prevent dehydration (Patterson Veterinary). A surgical drill (Model C300, Grobert) was used to make craniotomies for viral injections (all coordinates in reference to bregma). Craniotomies were made over the dorsal hippocampus (−2.1 mm anterior-posterior and −1.25 mm medial-lateral) or ventral hippocampus (−3.4 mm anterior-posterior, −2.7 mm medial-lateral). In a subset of experiments, a craniotomy was made over left dorsal CA3 (−2 mm anterior-posterior and −2.3 mm medial-lateral).

Viral labeling of MCs and MC axons was achieved using the Cre-dependent construct AAV5-EF1a-DIO-eYFP. Drd2-Cre mice were used to target either the dorsal or ventral hilus. Crlr-Cre mice were primarily used in experiments targeting the dorsal hilus because we observed that ventral hilar injections resulted in viral expression in a considerable number of CA3 pyramidal neurons, consistent with previous reports (Jinde et al., 2012; Yeh et al., 2018). Viral labeling of principal neurons in the dentate gyrus or hippocampal CA3 region was achieved using AAV5-CaMKIIa-ChR2(H134R)-mCherry.

Virus was injected using a 500 nL Neuros Syringe (#65457-02, Hamilton Company) attached to the stereotaxic apparatus. For MC targeting experiments, the syringe needle was slowly lowered into the craniotomy made over the dorsal hippocampus (1.9 mm below skull surface) or ventral hippocampus (3.4 mm below skull surface) and 150 nL of virus was injected at a rate of 80 nL/minute. In experiments targeting the hippocampal CA3 region, the needle was lowered mm below the skull surface and 100 nL of virus was injected at 80 nL/minute. In all experiments, the needle remained in place for at least 5 minutes after the injection to allow for diffusion of the virus before being slowly removed from the brain. The scalp was then cleaned with sterile saline and sutured using tissue adhesive (Vetbond, 3M). Mice were given 1 mL of lactated ringers (s.c.) at the end of surgery to support hydration. Mice were transferred to a clean cage at the end of the surgery and placed on a heating blanket (37 °C) until fully ambulatory.

### 2.3 Perfusions and sectioning

Mice were euthanized 14 days after surgery to evaluate viral expression. Mice were initially anesthetized with isoflurane, followed by urethane (2.5 g/kg; i.p.). Once under deep anesthesia, the abdominal cavity was opened and the subject was transcardially perfused with ∼10 mL of room temperature saline, followed by ∼20 mL of cold 4 % paraformaldehyde in 0.1 M phosphate buffer (PB; pH= 7.4). The brains were extracted and stored overnight at 4 °C in 4 % paraformaldehyde in 0.1 M PB. The brains were sectioned at 50 µm in the coronal or horizontal plane (Vibratome 3000, Ted Pella) and 1 of every 6 sections were selected for labeling (sections 300 µm apart). In a subset of experiments the left hemisphere was cut in the coronal plane, and the right hemisphere was cut in the horizontal plane to evaluate commissural projections of MCs. Sections were stored in 24-well tissue culture plates containing cryoprotectant (30 % sucrose, 30 % ethylene glycol in 0.1 M PB) at −20 °C until use (Botterill et al., 2015; Botterill et al., 2017).

### 2.4 Immunofluorescence

Immunofluorescence staining was performed on free floating sections as previously described (Botterill et al., 2019). A minimum of 5 sections per subject were used for immunofluorescence staining. Sections were washed in 0.1 M Tris Buffer (TB; 3 × 5 minutes each) and incubated in blocking solution consisting of 5 % normal goat serum, 0.25 % Triton X-100, and 1 % bovine serum albumin in 0.1 M TB for 30 minutes. To better visualize the MC axons, the viral label was amplified by incubating sections with chicken anti-GFP (1:2000, #ab13970, Abcam) or rabbit anti-mCherry (1:2000, #167453, Abcam) primary antibodies diluted in blocking solution.

For double labeling experiments, rabbit polyclonal anti-GluR2/3 (1:200, #AB1506, Millipore), mouse monoclonal anti-calretinin (1:750, #6B3, Swant), mouse monoclonal anti-GAD67 (1:500, #MAB5406, Millipore), or rabbit polyclonal vesicular GABA transporter (VGAT; 1:300, #131 003, Synaptic Systems) were added to the blocking solution containing primary antibodies against GFP and incubated overnight at 4 °C on a rotary shaker with gentle agitation **(Table 1)**. On the following day, the sections were washed in 0.1 M TB (3 ×5 minutes) and then incubated in goat anti-chicken Alexa 488 (1:1000, #A11039, Invitrogen), goat anti-rabbit Alexa 568 (1:500 to 1:1000, #A11036, Invitrogen), or goat anti-mouse Alexa 568 (1:500, #A11004, Invitrogen) secondary antibodies for 2 hours. The sections were then washed in 0.1 M TB (2 ×5 minutes) and counterstained with Hoechst 33342 (1:20000, #62249, Thermo Fisher Scientific) diluted in 0.1 M TB. The sections were then rinsed in 0.1 M TB (2 ×5 minutes), mounted onto gelatin-coated slides and air dried for 30 minutes. Sections were then coverslipped using Citifluor anti-fade mounting medium (#17970, Electron Microscopy Sciences).

**Table 1.** Antibody descriptions and parameters

| **Primary Antibodies** |  |  |  |  |  |  |
| --- | --- | --- | --- | --- | --- | --- |
| **Antigen** | **Host** | **Clonality** | **Dilution** | **Catalogue #** | **Vendor** | **RRID#** |
| GFP | Chicken | Polyclonal | 1:2000 | #AB13970 | Abcam | AB_300798 |
| mCherry | Rabbit | Polyclonal | 1:2000 | #167453 | Abcam | AB_2571870 |
| Calretinin | Mouse | Monoclonal | 1:750 | #6B3 | Swant | AB_10000320 |
| GAD67 | Mouse | Monoclonal | 1:500 | #MAB5406 | Millipore | AB_2278725 |
| VGAT | Rabbit | Polyclonal | 1:300 | #131 003 | Synaptic Systems | AB_887869 |
| GluR2/3 | Rabbit | Polyclonal | 1:100 | #AB1506 | Millipore | AB_90710 |
| **Secondary Antibodies** |  |  |  |  |  |  |
| **Antibody** | **Host** | **Visualization** | **Dilution** | **Catalogue #** | **Vendor** | **RRID#** |
| Alexa 488 Anti-Chicken | Goat | Fluorescence (488nm) | 1:1000 | A-11039 | Invitrogen | AB_142924 |
| Alexa 568 Anti-Rabbit | Goat | Fluorescence (568nm) | 1:500, 1:1000 mCherry | A-11036 | Invitrogen | AB_10563566 |
| Alexa 568 Anti-Mouse | Goat | Fluorescence (568nm) | 1:500 | A-11004 | Invitrogen | AB_2534072 |

### 2.5 Image acquisition

Images were acquired with a Zeiss LSM 880 laser scanning confocal microscope and Zen 3.0 software (Zeiss). Photomicrographs were acquired with Plan-Apochromat 10×/0,45 M27, Plan-Apochromat 20×/0.8 M27, or Plan-Apochromat 40x/1.4 Oil DIC M27 objectives. All images were acquired at 8-bit depth with a frame size of 1024×1024 or 2048×2048 pixels. For high-resolution insets, the Plan-Apochromat 40×/1.4 Oil DIC M27 objective was used with a 1.9 ×digital zoom. In cases where the region of interest was too large to fit within a single image (e.g., **Figure 1C7-C10**), tile scans were acquired with automatic stitching enabled in the acquisition software. Immunofluorescence was visualized with pre-configured excitation and emission wavelengths in the acquisition software for Hoechst 33342 (Ex/EM 408/453 nm), Alexa 488 / GFP (Ex/EM 488/535 nm), and Alexa 568 / mCherry (Ex/Em 561/643 nm). Zen 3.2 Blue Edition software (Zeiss) was used offline to export raw Zeiss image files (CZI format) into TIF format. Figures were made using Photoshop 21.2.3 (Adobe). When brightness and contrast adjustments were applied to a part of a figure, the same adjustments were made to each part of the figure.

**Figure 1.**
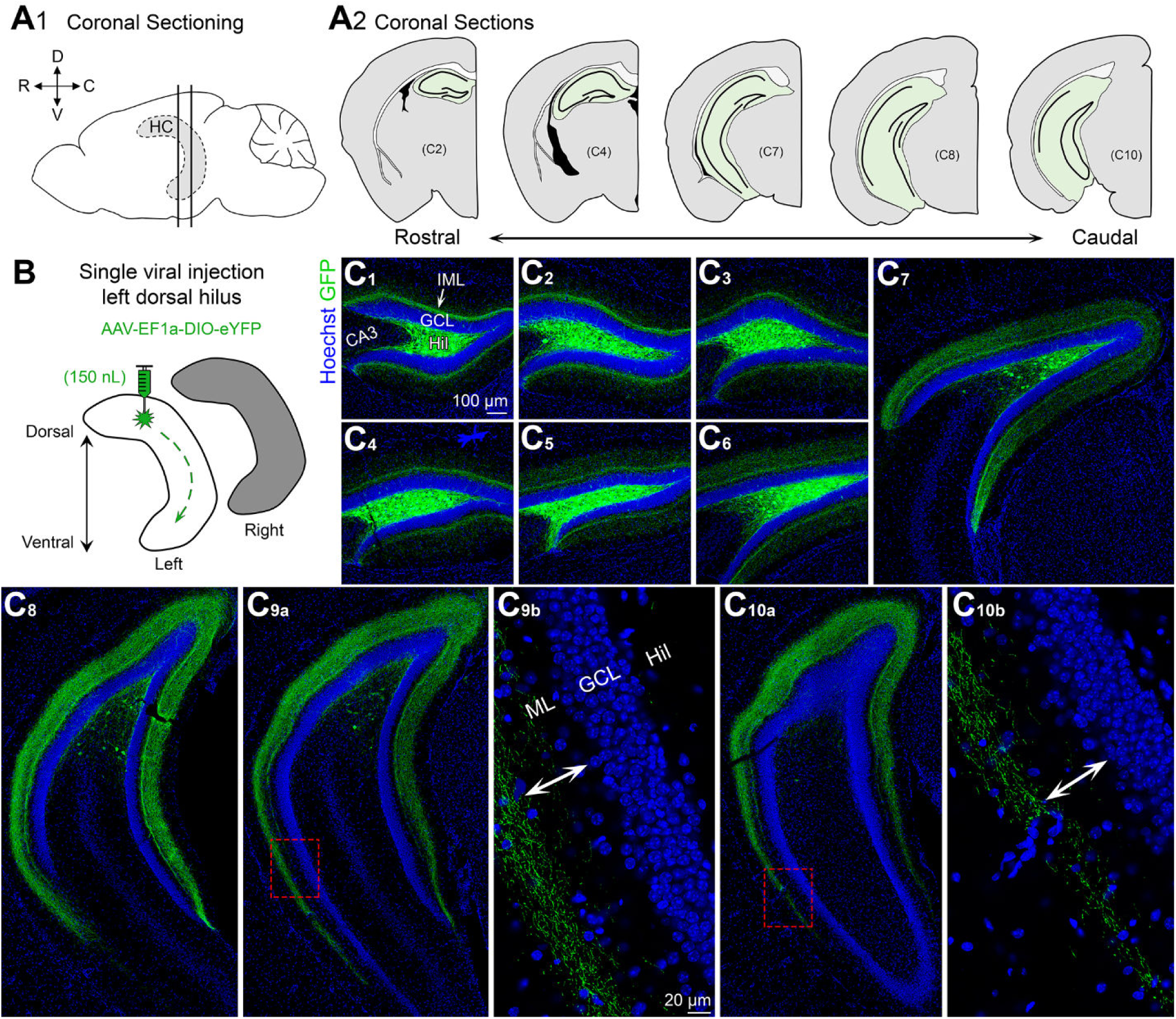
Viral expression of dorsal MCs and axons across the septotemporal axis of the DG. **(A1)** Side view of the brain showing the hippocampus (HC; grey with dashed border). Straight vertical lines are shown to depict sectioning in the coronal plane. (D) Dorsal (V) Ventral (R) Rostral (C) Caudal. **(A2)** Representative schematic of coronal sections starting from the rostral pole and extending to caudal hippocampus (green). **(B)** Viral injection schematic. 150nL of AAV-EF1a-DIO-eYFP was injected into the left dorsal hilus. The long-range axons of MCs are illustrated schematically with green dashes in the left hippocampus. Contralateral projections (right hippocampus; grey) are addressed in Figure 2. **(C)** Viral expression of MCs and their axons (green) across the septotemporal DG. Hoechst counterstain (blue) was used show the DG cell layer. **(C1-C4)** In the most dorsal hippocampus, GFP expression was primarily restricted to the hilus and the inner molecular layer (IML). **(C5-C8)** In progressively more caudal sections, fewer and fewer hilar GFP cells were observed, but the width of GFP axons in the molecular layer increased in the ventral parts of the sections. **(C9-C10)** In very caudal sections that included the most ventral parts of the DG, the GFP axons terminated in the middle to outer molecular layer in the ventral locations (arrows). (ML) molecular layer, (HIL) hilus, (GCL) granule cell layer.

### 2.6 Quantification of MC axons

Three measurements were made that are discussed and diagrammed in the Results and Figures. First, we measured the distance we call “inner”, corresponding to the gap that sometimes occurred between the GCL border with the IML and the GFP axon terminal plexus in the IML/MML. The gap was measured as the distance from the GCL border with the IML to the edge of the terminal plexus closest to the GCL. A schematic of measurements is shown in **Figure 5B3**. Next, we measured a distance we called “outer” which corresponded to the distance from the GCL border with the IML to the edge of the terminal plexus furthest from the GCL. Finally, a distance was measured called “width” which was defined as the distance between the edge closest and further from the GCL, i.e. the width of the GFP terminal plexus.

The distance measurements were made using the ‘distance tool’ in Zen 3.2 Blue. The length feature of the distance tool allows users to draw lines between two points to determine the distance between those points. These lines can be drawn in parallel, leading to the most precise measurements.

To define the GCL border with the IML, a line was drawn along the GCL border, defined by the Hoechst counterstain. The two edges of the GFP axon terminal plexus were defined readily because the plexus was a dense band of GFP puncta (reflecting MC axon boutons). Measurements were made for a region that was in the center of the upper blade, at the apex or crest of the DG, and the center of the lower blade to evaluate potential regional differences in viral expression.

Mice were injected in either the dorsal (n=6) or ventral (n=6) hilus and all analyses were done using horizontal sections because of their ability to clearly show the layers of the DG. In contrast, caudal DG in the coronal plane does not show the sublayers of ventral DG as well. A minimum of 3 dorsal and 3 ventral sections were analyzed for each subject.

### 2.7 Data analysis and statistics

All results are presented as the mean ± standard error of the mean (SEM). Statistical comparisons were made using Prism 8.4 (GraphPad) with statistical significance (*p*< 0.05) denoted on all graphs with an asterisk. Two-way ANOVAs were used for analyzing parametric data with multiple comparisons. Tukey’s post hoc test with corrections for multiple comparisons was used when appropriate.

## 3. RESULTS

### 3.1 GFP expression of MCs following a single dorsal hilus injection

#### 3.1.1 Coronal sections

Brains were sectioned in the coronal plane across the septotemporal axis of the DG **(Figure 1A1-A2)** to evaluate viral expression following a single injection into the left dorsal hilus **(Figure 1B)**. In dorsal sections proximal to the injection site **(Figure 1C1-C4)**, viral expression was observed strongly in the hilus and a weaker fluorescent signal was observed in the IML. Consistent with previous reports (Bernstein et al., 2020; Botterill et al., 2019), hilar GFP cell bodies near the injection site strongly colocalized with the glutamatergic marker GluR2/3 **(Figure S1)**. GFP axons in dorsal DG were primarily restricted to the IML **(Figure 1C1-C3)**. As sections progressed to more caudal regions of the hippocampus, the number of GFP cell bodies decreased significantly. However, the GFP axon became much wider and spread throughout the ML in caudal sections **(Figure 1C5-C8)**. The spread occurred in the part of the section that was more ventral. In extremely caudal sections, where the most ventral DG is visible, there were minimal hilar GFP cell bodies in the sections, but interestingly, the GFP axons in the ventral DG terminated broadly in both the MML and even the OML **(Figure 1C9-C10)**.

#### 3.1.2 Horizontal sections

As mentioned in the Methods, the hemisphere contralateral to the viral injection was sectioned horizontally. This allowed better evaluation of ventral hippocampus and also was used to examine the contralateral projection of MCs **(Figure 2A-B; S2)**. In contrast to past reports that the MC axon targets the contralateral DG in a homotopic fashion, we found that GFP axons were observed throughout the dorsal-ventral axis in the non-injected hemisphere. This finding suggests that commissurally-projecting MC axons are heterotopic and not homotopic as previously thought (Myers and Scharfman, 2009; Scharfman and Myers, 2012). In dorsal horizontal sections, GFP axons were observed throughout the ML **(Figure 2C1-2)**. As sections progressed from dorsal to more ventral hippocampus the GFP axon became increasingly further away from the GCL border **(Figure 2C3-C4)**. In the most ventral sections, the GFP axon was primarily in the MML with some labeling in the OML and almost no expression in the IML **(Figure 2C5-C6)**. Interestingly, scattered GFP hilar cells were observed throughout the dorsal-ventral axis of the contralateral (non-injected) DG **(Figure 2C1-C6)** although they were relatively rare compared to the dense labeling of somata at the injection site. High-resolution Z-stacks of the contralateral cells showed that they had morphology consistent with MCs, such as a large multipolar soma, numerous spiny dendrites, and dendritic regions with clusters of spines **(Figure S2)**. These contralateral cells may be a result of anterograde or retrograde labeling, which has been reported for multiple AAV serotypes, including AAV5 (Haery et al., 2019).

**Figure 2.**
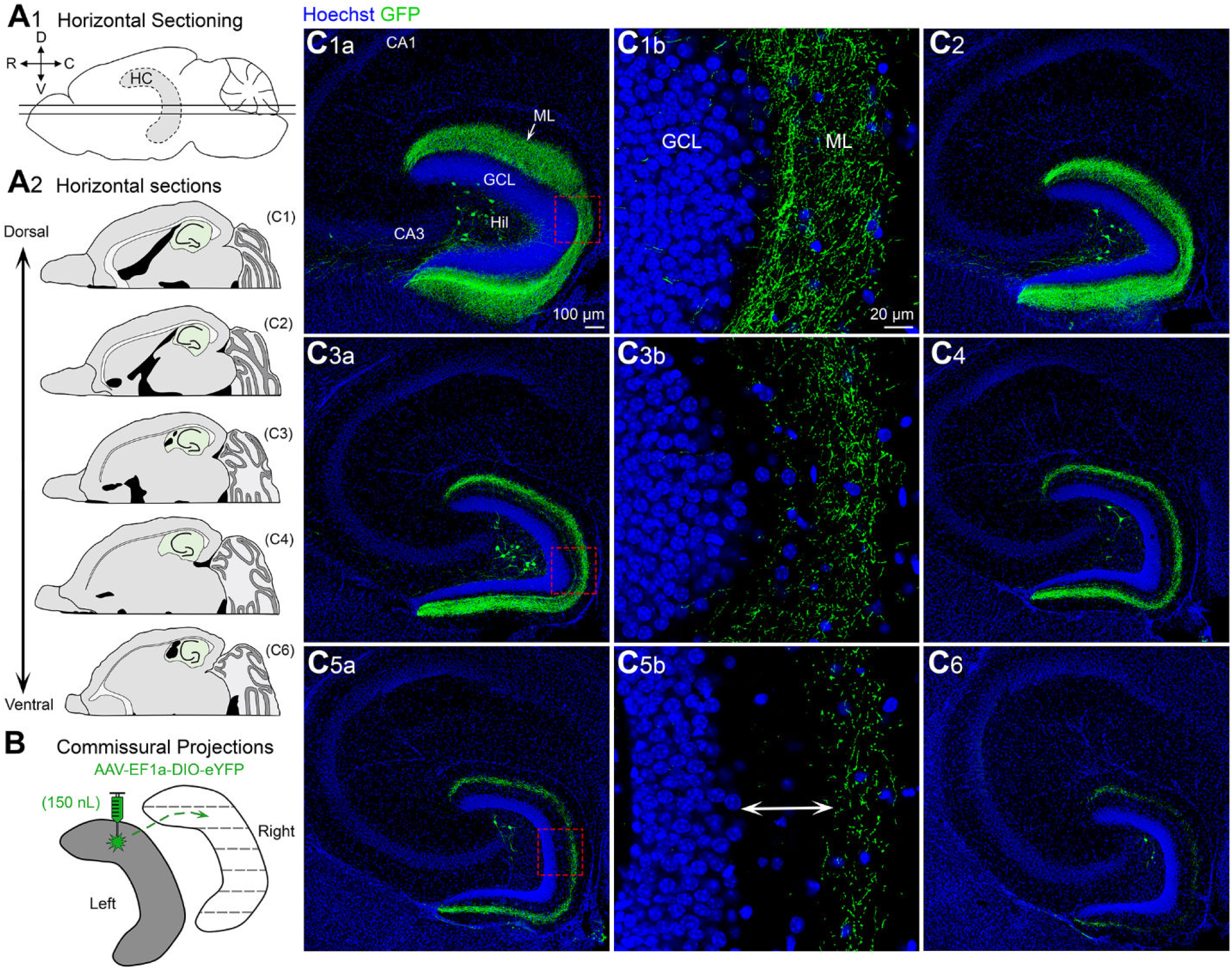
Contralateral projections of dorsal MC axons across the septotemporal axis of the DG. **(A1)** Side view of the brain showing the septotemporal extent of the hippocampus (HC; grey with dashed border). Straight horizontal lines are shown to illustrate the horizontal plane. (D) Dorsal (V) Ventral (R) Rostral (C) Caudal. **(A2)** Representative schematic of horizontal sections from a dorsal level to a progressively more ventral level (green). **(B)** To evaluate contralateral projections of dorsal MCs, the left hilus (grey) was injected and the right hippocampus (white) was evaluated in the horizontal plane. **(C)** Representative contralateral GFP axons are shown from sections that were dorsal and progressively more ventral. **(C1-C2)** In the relatively dorsal sections there were GFP axons throughout the molecular layer. **(C3-C4)** “Mid” sections (between the dorsal sections in C1-C2 and the ventral sections in C5-C6) showed GFP axons that terminated increasingly further away from the GCL border. **(C5-C6)** GFP axons in ventral sections were in the MML/OML primarily (arrow). (ML) molecular layer, (HIL) hilus, (GCL) granule cell layer.

### 3.2 GFP expression following a single ventral hilus injection

#### 3.2.1 Coronal sections

In a separate set of experiments, mice received a single viral injection into the left ventral hilus **(Figure 3A)**. Similar to the dorsal injection, a single ventral injection also resulted in GFP axon labeling throughout the entire septotemporal extent of the DG **(Figure 3B)**. In dorsal sections, i.e., distal to the injection site, there were no GFP cells within the hilus **(Figure 3B1-B4)**. As sections progressed to more caudal and ventral regions, the number of GFP hilar cells increased significantly **(Figure 3B5-B10)**. A very small number of weakly-labeled GFP cells were observed in the CA3c region of some sections **(Figure 3B7-B9)**, consistent with previous reports (Fredes et al., 2019; Yeh et al., 2018). Importantly, the GFP axon was largely restricted to the IML of the DG throughout the entire septotemporal axis of the DG. This result suggests that dorsal and ventral MCs have distinct axonal projections.

**Figure 3.**
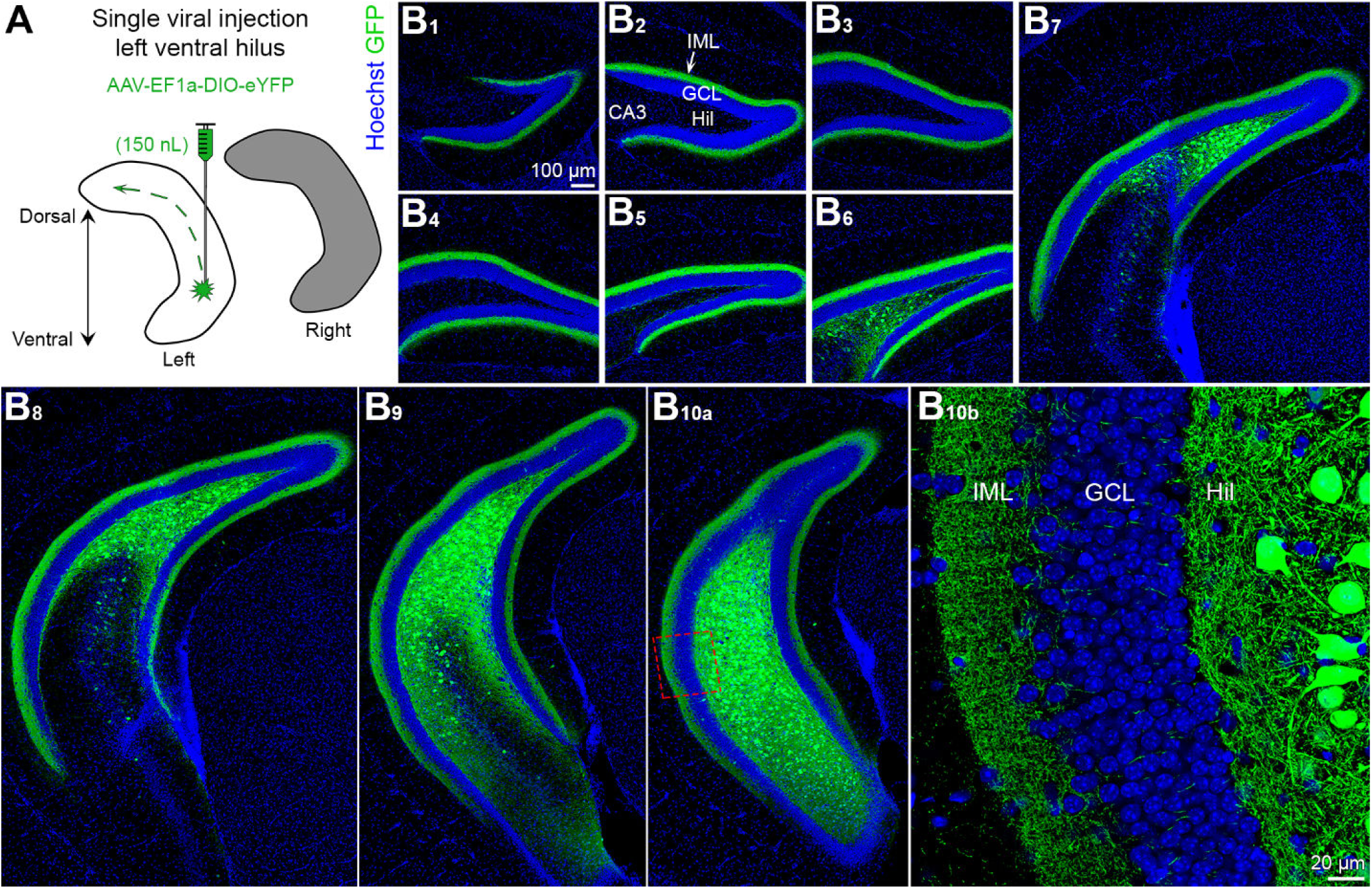
Viral expression in ventral MCs and their axons across the septotemporal axis of the DG. **(A)** Viral injection schematic. AAV-EF1a-DIO-eYFP was injected into the left ventral hilus. The long-range axons of ventral MCs are depicted with the green dashes in the left hippocampus (white). Contralateral projections of ventral MCs (right hippocampus; grey) are addressed in Figure 4. **(B)** Viral expression of ventral MCs and their axons (green) across the septotemporal DG. Hoechst counterstain (blue) was used show the DG cell layer. **(B1-B5)** In the dorsal hippocampus, GFP expression was primarily restricted to the inner molecular layer (IML). **(B6-B8)** In sections that were progressively more caudal, GFP expression was observed in the hilus and IML in the part of the DG that was more ventral. **(B9-B10)** In sections that included the most ventral part of the DG sections, GFP expression in ventral locations was observed in the hilus and IML. (IML) inner molecular layer, (HIL) hilus, (GCL) granule cell layer.

#### 3.2.2 Horizontal sections

To best evaluate the commissural projections of ventral MCs throughout the dorsal-ventral axis, brains were hemisected and the right (non-injected) hemisphere was cut in the horizontal plane **(Figure 4A)**. Similar to dorsal hilar injections, mice with a ventral hilar injection showed GFP axon expression throughout the entire dorsal-ventral axis of the contralateral DG **(Figure 4B)**. This observation provides further support for the notion that contralateral MC axons are heterotopic and not homotopic (as discussed above). Furthermore, similar to the coronal sections, the GFP axon was restricted primarily to the IML throughout the entire dorsal-ventral axis **(Figures 2B1b, 2B3b, 2B5b)**. Interestingly, unlike dorsal injections, mice injected in the ventral hilus had few or no GFP cells in the hilus of the contralateral DG.

**Figure 4.**
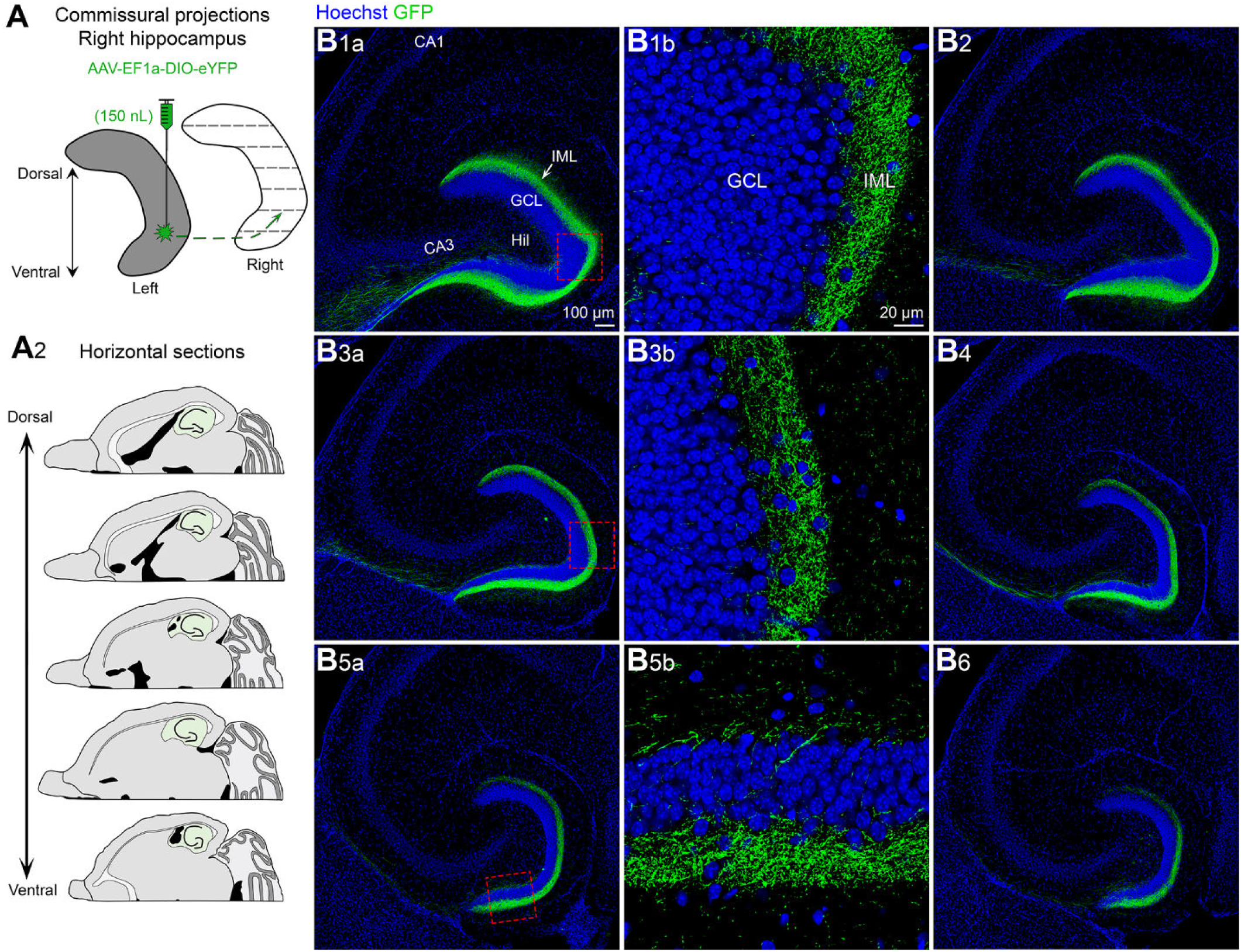
Contralateral projections of ventral MC axons across the septotemporal axis of the DG. **(A1)** To evaluate contralateral projections of ventral MCs, the left hilus was injected with AAV-EF1a-DIO-eYFP (grey) and the right hippocampus (white) was evaluated in the horizontal plane. **(A2)** Representative schematic of horizontal sections from dorsal to more ventral hippocampus (green). **(B)** Representative examples of GFP axons from dorsal levels to progressively more ventral locations. **(B1-B6)** The contralateral projections of ventral MCs appear to be primarily restricted to the IML in all sections (IML) inner molecular layer, (HIL) hilus, (GCL) granule cell layer.

**Figure 5.**
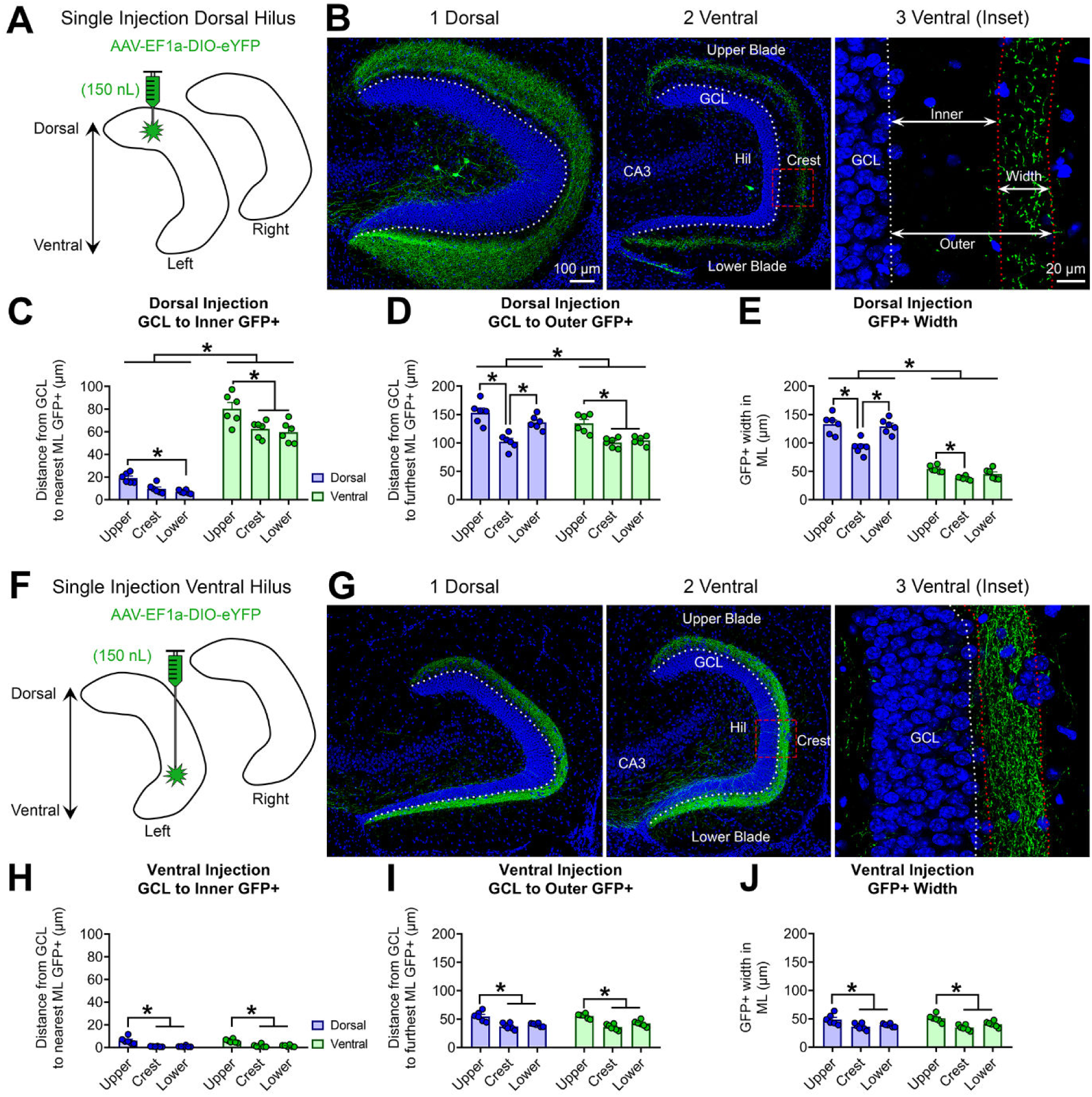
Quantitative analysis of the GFP MC axon terminal plexus. **(A)** Schematic showing that AAV-EF1a-DIO-eYFP was injected into the left dorsal hilus. **(B1-B2)** Representative expression of contralateral GFP axon terminals in the **(B1)** dorsal and **(B2)** ventral dentate gyrus. **(B3)** A schematic shows the inner, outer, and width measurements for the GFP axon plexus. Measurements were made in the center of the upper blade, crest and center of the lower blade.**(C)** The distance between the GCL and closest part of the GFP axon terminal field (“inner”) was significantly greater in ventral sections relative to dorsal sections. **(D)** The distance between the GCL and furthest border of the GFP axon terminal field from the GCL (“outer”) was significantly greater in dorsal relative to ventral sections. **(E)** The total width of the GFP axon plexus (“width”) was significantly greater in dorsal relative to ventral sections. **(F)** A schematic for additional animals where AAV-EF1a-DIO-eYFP was injected into the left ventral hilus. **(G1-G2)** Representative expression of contralateral GFP axons in the relatively **(G1)** dorsal and **(G2)** ventral dentate gyrus. **(G3)** A schematic showing GFP axon measurements which were the same as above. **(H)** The inner distance did not differ between dorsal and ventral sections. **(I)** The outer distance did not differ between dorsal and ventral sections. **(J)** The width did not differ between dorsal and ventral sections. \**p*<0.05.

### 3.3 Measurements of the GFP axon in dorsal and ventral injected mice

Next, we sought to quantify the previously described differences in GFP axonal expression following a single dorsal (n=6) or ventral (n=6) hilar injection.

#### 3.3.1 GFP axon measurements for dorsal viral injections

Following a single dorsal viral injection **(Figure 5A)**, we evaluated GFP axon distance in dorsal and ventral sections **(Figure 5B1-2)**. Using the GCL border with the IML (GCL/IML border or GCL “outer” border below) as a reference point, we measured the distances defined as inner, outer, and width in the Methods and shown in **Figure 5B3**. First, we measured the distance from the GCL/IML border to the start of the band of GFP immunofluorescence in the IML/MML **(Figure 5C)**. A two-way ANOVA revealed a significant main effect of septotemporal location (*F*(1,30)=468.0, *p*<0.001), attributable to dorsal sections (11.89 ± 1.49 µm) having a shorter GFP-IML distance than ventral sections (67.50 ± 3.19 µm). Thus, there was little gap between the GCL and MC axons dorsally but ventrally there was a notable gap.

We also observed a main effect of upper vs. lower blade (*F*(2,30)=15.68, *p*<0.001). In dorsal sections, with the distance significantly greater in the upper blade (19.11 ± 1.78 µm) compared to the lower blade (6.91 ± 0.65 µm; *p*=0.027). In ventral sections, the distance was significantly greater in the upper blade (80.39 ± 5.45 µm) compared to the crest (62.51 ± 2.98 µm) and lower blade (59.61 ± 3.81 µm; all *p* values <0.01).

Next, we measured the distance from the GCL/IML border to the point where the GFP terminal plexus ended either along the IML/MML border, or in the MML/OML **(Figure 5D)**. A two-way ANOVA revealed a significant main effect of septotemporal location (*F*(1,30)=14.93, *p*<0.001), attributable to the GFP terminal plexus reaching more of the MML and even OML in dorsal (130.8 ± 6.16 µm) compared to ventral sections (113.4 ± 4.46 µm). We also observed a main effect of blade (*F*(2,30)=29.66, *p*<0.001), and a significant interaction (*F*(2,30)=3.762, *p*=0.034). Tukey’s post hoc test revealed that the distance was significantly greater in the upper (153.6 ± 7.89 µm) and lower blades (136.5 ± 4.91 µm) compared to the crest (102.5 ± 5.48 µm) in dorsal sections (all *p* values <0.001). In ventral sections, the distance was significantly greater in the upper blade (134.7 ± 6.64 µm) than the crest (100.9 ± 3.54 µm) and lower blade (104.7 ± 3.12 µm; all *p* values <0.001).

We also measured the total width of the GFP terminal plexus in the ML **(Figure 5E)**. A two-way ANOVA revealed a significant main effect of septotemporal location (*F*(1,30)=373.7, *p*<0.001), with dorsal sections (118.5 ± 5.46 µm) having a significantly wider GFP axon than the ventral sections (45.84 ± 2.17 µm). The results also revealed a main effect of blade (*F*(2,30)=20.73, *p*<0.001), and a significant interaction (*F*(2,30)=5.943, *p<*0.001. In dorsal sections, the GFP axon was significantly wider in the upper (133.1 ± 7.11 µm) and lower blades (129.6 ± 5.27 µm) compared to the crest (92.76 ± 5.14 µm; all *p* values <0.001). In ventral sections, the GFP axon was significantly wider in the upper blade (54.29 ± 2.37 µm) compared to the crest (38.14 ± 1.25 µm; *p*=0.048).

#### 3.3.2 GFP axon measurements following a ventral viral injection

Using the same approach as above, we quantified sections from mice injected in the ventral hilus **(Figure 5F-G)**. First, regarding the “gap” between the GCL and the GFP terminal plexus **(Figure 5H)**, a two-way ANOVA revealed a significant main effect of blade (*F*(2,30)=38.63 *p*<0.001) but no differences between septotemporal locations (*F*(1,30)=0.216, *p*=0.645). Within dorsal sections, the distance was significantly greater in the upper blade (6.03 ± 1.21 µm) compared to the crest (0.77 ± 0.10 µm) and lower blade (0.85 ± 0.25 µm; all *p* values <0.001). Similarly, within ventral sections, the distance was significantly greater in the upper blade (5.64 ± 0.64 µm) compared to the crest (1.36 ± 0.52 µm) and lower blade (1.37 ± 0.30 µm; all *p* values <0.001).

Next, for the distance from the GCL/IML border to the end of the GFP immunofluorescence in the outer portion of the ML **(Figure 5I), a** two-way ANOVA revealed a significant main effect of blade (*F*(2,30)=35.26 *p*<0.001) but no difference in septotemporal location (*F*(1,30)=0.070, *p*=0.792). Tukey’s post hoc test revealed that the distance was significantly greater in the upper blade (54.81 ± 3.47 µm) than the crest (37.21 ± 2.35 µm) and lower blade (40.14 ± 1.22 µm; all *p* values <0.001) in dorsal sections. We observed a similar result in ventral sections, with the distance being greater in the upper blade (55.33 ± 1.72 µm) than the crest (36.03 ± 2.16 µm) or lower blade (42.28 ± 2.15 µm; all *p* values <0.001)

For the width of the GFP immunofluorescence in the ML **(Figure 5J)**, a two-way ANOVA revealed a significant main effect of blade (*F*(2,30)=17.27 *p*<0.001) but no difference between septotemporal location (*F*(1,30)=0.085, *p*=0.772). Tukey’s post hoc test showed that in dorsal sections, the width was significantly greater in the upper blade (48.67 ± 3.76 µm) than the crest (36.13 ± 2.40 µm) or lower blade (38.89 ± 1.18 µm; all *p* values <0.025). Similarly, in ventral sections the width was significantly greater in the upper blade (50.58 ± 2.80 µm) than the crest (34.29 ± 2.02 µm) or lower blade (40.60 ± 2.10 µm; all *p* values <0.022).

### 3.4 CaMKIIa injections in the hilus result in a similar pattern of long-range and commissural ML expression as MC-specific targeting

Next, we used a different approach than Drd2 or Crlr-Cre mice because of the possibility that these mouse lines express virus in hilar GABAergic neurons. To this end, we targeted excitatory neurons in the DG using a viral construct that utilized the calcium/calmodulin-dependent protein kinase II (CaMKIIa) promoter. A virus using a mCherry tag was used instead of GFP simply due to availability of viruses. This approach also labels excitatory cells like the GCs and CA3c pyramidal neurons, but this was actually useful as explained below.

#### 3.4.1 CaMKIIa-mCherry injection into the dorsal hilus

The dorsal hilus was injected with AAV-CaMKIIa-ChR2(H134R)-mCherry using identical parameters as the Cre-dependent expression experiments **(Figure 6A)**. DrD2-Cre^-/-^ mice (n=3) were used since the Cre^+/-^ mice were not needed for viral expression and not valuable in these experiments, as explained above. Brains were sectioned in the horizontal plane. In sections near the injection site, mCherry viral expression was observed in GCs, mossy fibers, hilar cells (putative MCs), and CA3 pyramidal neurons **(Figure 6B1-B2)**, consistent with the selectivity of CaMKIIa for excitatory neurons. As sections were evaluated in more ventral regions, mCherry expression in the ML became increasingly further from the GCL, consistent with the pattern observed when MCs were targeted selectively **(Figure 6B3-B7)**. In addition, mCherry expression was also observed in the CA3 stratum radiatum of all sections, presumably due to targeting of the Schaffer collateral axons of CA3 pyramidal neurons.

**Figure 6.**
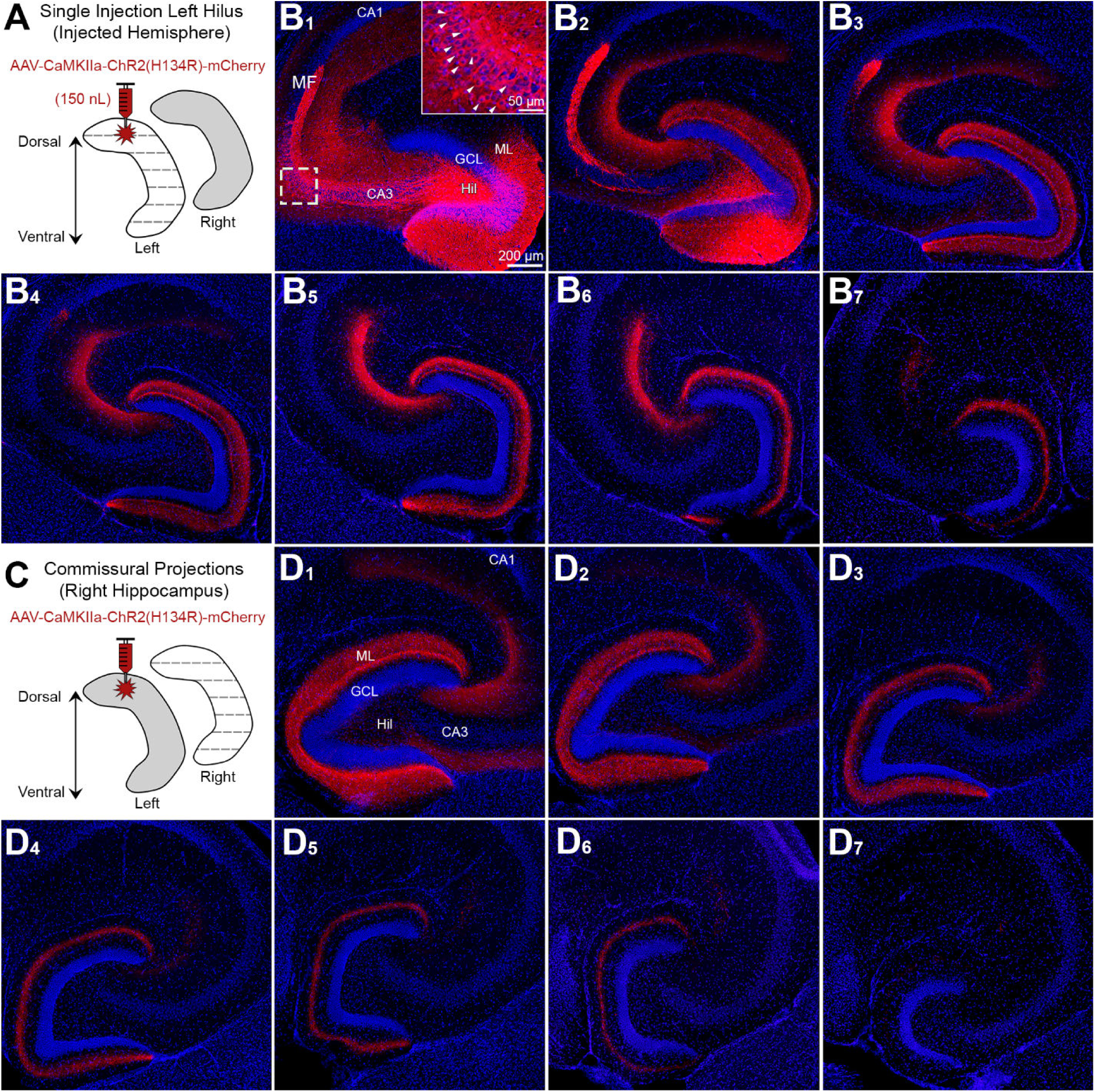
Use of CaMKIIa to probe the specificity of GFP for MCs. **(A)** Viral injection schematic. 150nL of AAV-CaMKIIa-ChR2(H134R)-mCherry was injected into the left dorsal hilus to target excitatory neurons. **(B1-B2)** Proximal to the injection site, viral expression was observed in GCs, MCs, and CA3 pyramidal neurons (inset; white arrowheads). Granule cell mossy fibers (MF) axons were also labeled where they normally project, CA3 stratum lucidum. **(B3-B7)** Long-range mCherry axons showed a similar pattern of viral expression in the molecular layer as Drd2-Cre or Crlr-Cre mice injected in the dorsal DG with a virus to express GFP. **(C)** Contralateral axons were evaluated in the right hippocampus. **(D1-D7)** Contralateral mCherry axons showed a similar pattern in the molecular layer as Drd2-Cre and Crlr-Cre injected with a virus expressing GFP in the dorsal hilus. This figure shows that injection of AAV to express CaMKIIa in the dorsal hilus results in a similar pattern of axon labeling as an injection of AAV to express GFP in MCs. (ML) molecular layer, (HIL) hilus, (GCL) granule cell layer.

Commissural projections were also assessed by evaluating the non-injected hemisphere of the same mice **(Figure 6C)**. Consistent with our previous experiments, a similar pattern of mCherry expression was observed across the dorsal-ventral axis, whereby mCherry expression was seen throughout the ML in dorsal sections (e.g., **Figure 6D1**) and selective to the MML-OML of ventral sections (e.g., **Figure 6D4-D6**). Interestingly, contralateral mCherry expression was also observed in the CA3 stratum radiatum of dorsal sections (**Figure 6D1-D3**), which was not seen in experiments targeting the MCs only.

Taken together, mCherry expression in the ipsilateral and contralateral ML was similar to MC-specific experiments that targeted the dorsal hilus. These results support the notion that the dorsal-ventral distribution of GFP axons described in previous experiments are attributable to MCs rather than non-specific targeting of GABAergic hilar cell populations. They also support the idea that GCs and pyramidal neurons of CA3 did not contribute significantly to data using GFP in Drd2-Cre or Crlr-Cre mice, and the role of CA3 is addressed further below.

#### 3.4.2 CaMKIIa-mCherry injection into the dorsal CA3 region

Given the observation that CA3 neurons can be labeled in Drd2-Cre or Crlr-Cre lines, we targeted the CA3 area with virus to determine whether viral expression in CA3 can contribute to viral expression in the ML. Drd2-Cre^-/-^ (n=3) mice were injected in the dorsal CA3 (a/b subfield) with AAV-CaMKIIa-ChR2(H134R)-mCherry **(Figure S3A)**. Pilot experiments found that larger volumes or injections more proximal to CA3c labeled MCs and therefore prevented us from determining whether CA3 could contribute to ML immunofluorescence. In sections proximal to the injection site, we observed viral expression in the CA3 pyramidal cell layer **(Figure S3B1-B3)**. We also observed a band of mCherry expression in CA3 stratum radiatum, supporting the notion that CA3 pyramidal neurons caused the stratum radiatum mCherry expression in the CaMKIIa experiments that targeted the hilus **(Figure 6)**. Importantly, throughout the dorsal-ventral axis, there was no mCherry immunofluorescence in the ML. Taken together, these results suggest that the long-range mCherry axons in the ML of hilar-injected mice were due to MCs and not CA3 pyramidal neurons.

### 3.5 Ventral but not dorsal MCs correspond to calretinin immunoreactivity

In the mouse, calretinin is widely used as a marker for MC somata and MC axons in the IML. Consistent with past reports (Blasco-Ibanez and Freund, 1997; Fujise et al., 1998), calretinin expression of MC somata is primarily observed in the ventral hilus but not dorsal hilus **(Figure S4)**. However, calretinin immunoreactivity in the IML is observed throughout the entire septotemporal axis of the DG (Blasco-Ibanez and Freund, 1997; Fujise et al., 1998). This led us to hypothesize that calretinin IML immunoreactivity is due to ventral but not dorsal MCs.

First, we evaluated mice injected in the dorsal hilus with AAV-EF1a-DIO-eYFP and sections were processed for calretinin immunofluorescence (n=3; **Figure 7A**). In dorsal sections **(Figure 7B1)**, we found that calretinin immunofluorescence was observed in the IML; however, cell bodies in the hilus were not labeled with calretinin. In contrast, GFP expression was strongly expressed in hilar cells and moderately expressed in the IML, resulting in minimal colocalization of calretinin and GFP **(Figure 7B2-7B4)**. In ventral sections, calretinin immunofluorescence was observed in hilar cells and the IML **(Figure 7B5-B6)**. Remarkably, GFP axons terminated in the MML-OML, adjacent to the calretinin immunofluorescence in the IML **(Figure 7B7-B8)**. This result is consistent with prior studies showing that dorsal MC somata lack calretinin expression. It also helps explain why the MML-OML projection of dorsal MCs has not been reported using classic immunohistochemical approaches. Indeed, it appears that viral labeling is required to study dorsal MCs and their unique long-range axons.

**Figure 7.**
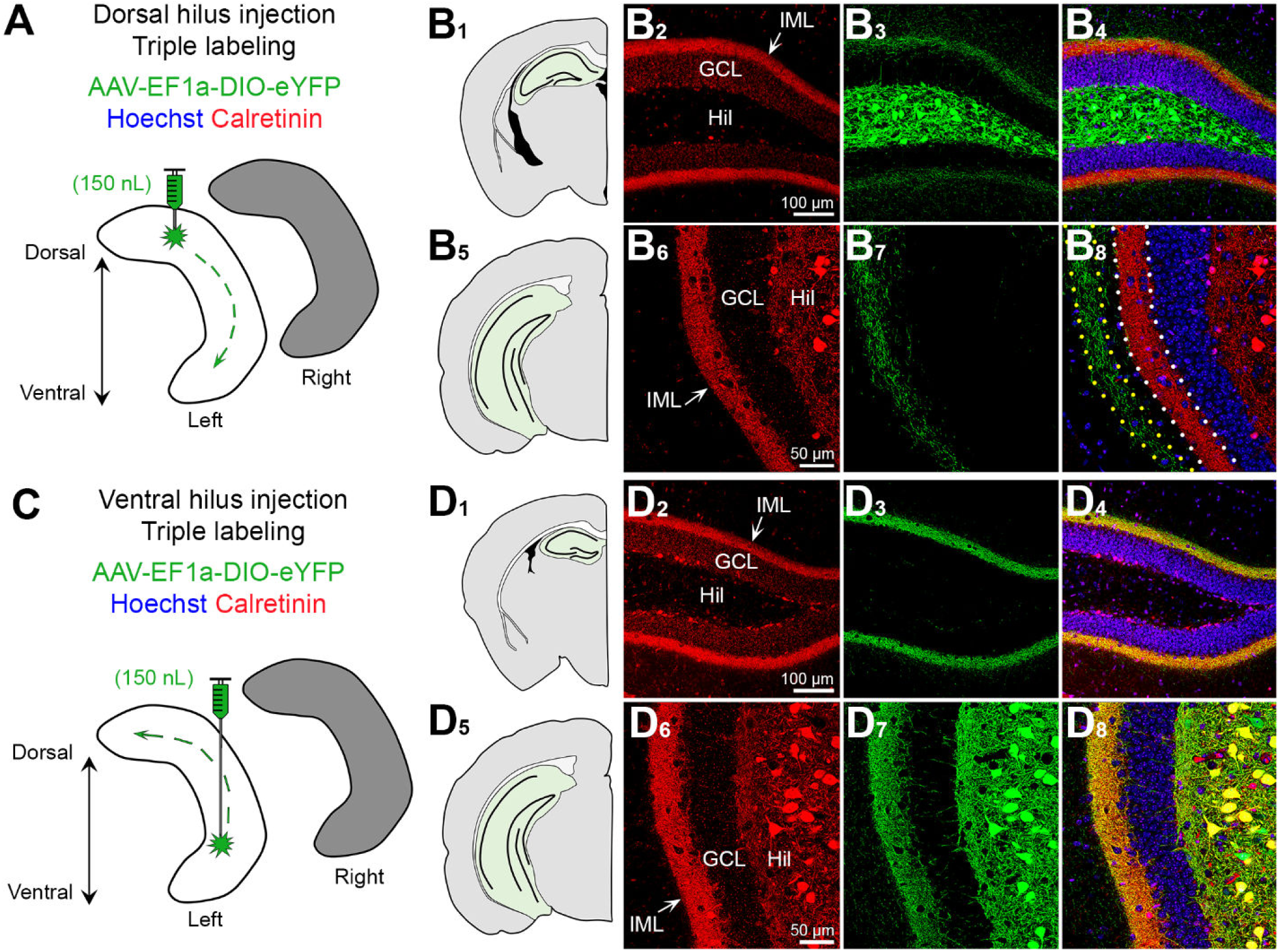
Calretinin labels ventral but not dorsal MCs. **(A)** Viral injection schematic. AAV-EF1a-DIO-eYFP was injected into the left dorsal hilus. **(B1-B4)** In the dorsal DG, calretinin (red) is primarily in the IML of the DG, whereas viral expression (green) is strong in hilar cells and weak in the IML. **(B5-B8)** In ventral hippocampus, calretinin expression (red) is in putative hilar MCs and the IML. Long-range viral-expressing axons (green) are observed in the molecular layer adjacent to calretinin immunofluorescence in the IML (dotted borders). **(C)** Viral injection schematic. AAV-EF1a-DIO-eYFP was injected into the left ventral hilus. **(D1-D4)** Calretinin (red) and GFP long-range axons are primarily in the IML and appear to colocalize (yellow). **(D5-D8)** In ventral sections, calretinin (red) and GFP strongly overlap within hilar cell bodies and the IML (yellow). (IML) inner molecular layer, (HIL) hilus, (GCL) granule cell layer.

Next, we evaluated mice (n=3) injected in the ventral hilus with AAV-EF1a-DIO-eYFP and processed sections for calretinin immunofluorescence **(Figure 7C)**. In dorsal sections **(Figure 7D1)** we found that calretinin and GFP immunofluorescence were primarily in the IML and showed strong colocalization **(Figure 7D2-D4)**. In ventral sections **(Figure 7D5)** we found that calretinin and GFP immunofluorescence was similar and showed a high degree of colocalization in the IML (MC axons) and hilus (cell bodies; **Figure 7D6-D8)**. These results suggest that ventral MCs express calretinin in their cell bodies and their long-range axons in the IML across the dorsal-ventral axis of the DG.

### 3.6 GFP axons in the ML show minimal colocalization with GABAergic markers

Next, we sought to determine whether non-specific targeting of GABAergic neuron axons contributed to viral expression in the ML. Notably, GABAergic hilar neurons such as hilar perforant path-associated (HIPP) cells are hilar cells with axons that project locally to the hilus, MML-OML and to the contralateral MML-OML (Deller and Leranth, 1990; Eyre and Bartos, 2019). To address the potential concern that some of the GFP axons were due to HIPP cells, we injected mice in either the dorsal (n=3) or ventral (n=3) hilus with AAV-EF1a-DIO-eYFP and processed the tissue with two widely used antibodies for GABAergic terminals: VGAT and GAD67. By using these two markers, we also could address the possibility that some of the axons in the IML were due to HICAP cells (Halasy and Somogyi, 1993; Han et al., 1993), and some axons in the MML or OML were from MOPP cells (Halasy and Somogyi, 1993; Han et al., 1993) or molecular layer neurogliaform cells (Armstrong et al., 2011).

#### 3.6.1 VGAT

First, we evaluated VGAT immunofluorescence in mice injected in the left dorsal hilus with AAV-EF1a-DIO-eYFP **(Figure S5A)**. VGAT immunofluorescent terminals were observed around GC somata and throughout the ML **(Figure S5B**), consistent with previous studies of GABAergic terminal distribution in the DG (Freund and Buzsaki, 1996; Houser, 2007). In both dorsal and ventral sections, the GFP axon in the ML failed to show clear colocalization with VGAT **(Figure S5B1-B2)**. However, GFP terminals were often adjacent to or near VGAT+ puncta, which is not surprising given the density of MC and GABAergic labeling. In a few cases GFP and VGAT+ immunofluorescence appeared to overlap and produce a yellow product, but this was due to a GFP bouton on or overlapping a GABAergic bouton in a different focal plane. We also evaluated VGAT immunofluorescence in mice that received AAV-EF1a-DIO-eYFP in the left ventral hilus **(Figure S5C)**. In mice injected in the ventral hilus, the GFP axons were primarily restricted to the IML **(Figure S5D1-D2)**. Similar to the dorsally-injected mice, the GFP axons showed minimal colocalization with VGAT in both dorsal and ventral sections. Taken together, these results suggest that the GFP axons in dorsally- and ventrally-injected mice were unlikely to be due to GABAergic terminals. This finding is further supported by the CaMKIIa experiments that targeted excitatory neurons in the dorsal hilus that produced a similar pattern of ML immunofluorescence across the dorsal-ventral axis as GFP.

#### 3.6.2 GAD67

Next, we evaluated GAD67 immunofluorescence in mice that received a viral injection of AAV-EF1a-DIO-eYFP into the dorsal or ventral hilus **(Figure S6A & S6C)**. Similar to VGAT, GAD67 was observed around GCs and throughout the ML; however, GAD67 also resulted in some somatic labeling throughout the hilus and ML **(Figure S6B1 & S6D2)**. In mice where the viral injection was the dorsal hilus, we observed minimal GFP and GAD67+ colocalization and this was true for sections that were located throughout the dorsal-ventral axis **(Figure S6B1-B2)**. A similar observation was made for mice that received viral injections in the ventral hilus. Indeed, both dorsal and ventral sections showed minimal GFP/GAD67+ colocalization **(Figure 6D1-D2)**. In summary, GAD67 immunofluorescence showed minimal colocalization in GFP axons, suggesting that the GFP axons are primarily GABA negative. This finding is further supported by the VGAT immunofluorescence which also showed minimal colocalization with GFP immunofluorescent MC axons.

### 3.7 Mistargeted viral injections do not cause ML GFP expression

Finally, we show that GFP expression is absent in the ML of animals that received viral injections that were outside of the DG. These injections were accidental and due to experimenter error such as misreading the coordinates of the stereotaxic apparatus, head tilts, or lowering the injection syringe to an inaccurate depth (or any combination of these factors). In one representative example, we found that an accidental injection in the thalamus of a Drd2-Cre^+/-^ resulted in GFP cell expression proximal to the injection site, but no expression was observed in the DG **(Figure S7)**. Thus, when virus labeled areas surrounding but not within DG, we observed no viral expression in the DG. Taken together with the previous results, these data suggest that viral expression in the ML required viral expression in hilar cells and did not arise from other local sources (e.g., CA3; **Figure S3**) or regions outside of the DG such as the thalamus.

## 4. DISCUSSION

### 4.1.1 Differences between dorsal and ventral MCs

The results showed significant differences in the axonal projections of dorsal and ventral MCs. This is important because most investigators currently consider MCs to be a homogeneous population. In the past, there have been a few published papers where differences between dorsal and ventral MCs have been reported but they are rare. Therefore, our demonstration of significant differences in dorsal and ventral mouse MCs could have an impact on future investigations.

The past studies showing dorsal-ventral differences in MCs are mainly in the rat. For example, it has been shown that calretinin expression is high in ventral MC somata but not dorsal MCs (Freund and Buzsaki, 1996; Kosaka et al., 1987) a result we replicated in the present study. Another study which suggested that dorsal and ventral MCs were different was electrophysiological, and used hippocampal slices to show that ventral MCs exhibited a greater degree of bursts in response to pharmacological agents (Jinno et al., 2003). More recently, a study in transgenic mice showed that ventral MCs have significantly different effects on behavior compared to dorsal MCs (Fredes et al., 2019).

The differences in dorsal vs. ventral MCs are important because they may contribute to the dorsal and ventral differences in DG function that have been widely discussed (Chawla et al., 2018; Kheirbek et al., 2013; Kheirbek and Hen, 2011). The MC axon could play a role in these dorsal-ventral differences because dorsal MCs project primarily to ventral locations in the ipsilateral hippocampus and the opposite is true for ventral MCs. Thus, ventral MCs primarily innervate dorsal GCs in the ipsilateral hippocampus. If a broader terminal plexus leads to different effects than a restricted plexus, which seems like a reasonable prediction, dorsal MCs would influence dorsal GCs differently than ventral GCs. In contrast, ventral MCs will have similar effects on GCs, regardless of the dorsal or ventral GC location. Therefore, dorsal MCs may differentially affect GCs whereas ventral MCs may have a more homogeneous effect.

### 4.1.2 Dorsal MC axons in the IML expand to include the MML in ventral and contralateral DG

The experimental data used many approaches to confirm the results. For example, two different transgenic mouse lines with Cre recombinase expressed in MCs were used. This study provides several lines of evidence that dorsal MCs have an axon restricted to the IML in dorsal sites near the MC cell body. In contrast, the axon terminates primarily in the MML of distal sites in the ventral and contralateral DG. In contrast, ventral MCs did not share these characteristics, only showing terminations in the IML throughout the septotemporal axis.

There also might be contamination by GABAergic neurons of the DG that project to the MML, which have axons that collectively cover the GC somatodendritic axis (Freund and Buzsaki, 1996; Houser, 2007). However, there is no type of DG GABAergic neuron that projects only to the MML. Hilar GABAergic neurons which express somatostatin and NPY do have projections to the molecular layer, but their axons are distributed to the outer two-thirds, not the middle third (Deller and Leranth, 1990; Eyre and Bartos, 2019; Freund and Buzsaki, 1996; Houser, 2007; Sperk et al., 2007). Molecular layer GABAergic neurons such as MOPP cells (Halasy and Somogyi, 1993) or neurogliaform cells (Armstrong et al., 2011) may have an axon that is restricted to the molecular layer but there are several characteristics about the axons of these GABAergic neurons that are different from the axonal distribution we observed in the MML. What we found was GFP axonal terminals throughout the MML were robust throughout the lateral tip of the upper blade all the way around the DG to the lateral tip of the lower blade. In other words, a homogeneous band of fibers stained the MML throughout the DG in any given section. In contrast, the MOPP cell and neurogliaform cells have an axon that is localized to the area around their somata and this includes both the OML and MML (Armstrong et al., 2011; Halasy and Somogyi, 1993). Notably, the Drd2-Cre mouse has been suggested to show expression of Cre not only in MCs but also some hippocampal GABAergic neurons (Puighermanal et al., 2015), but we have found this rare (Bernstein et al., 2020; Botterill et al., 2019). Nevertheless, in the present paper we used two markers of GABAergic neurons and asked if there was colocalization of viral expression of MCs with GABAergic neuron labeling. The results did not show evidence of double-labeling, making it unlikely that there was significant contamination of GFP expression by GABAergic neurons.

In Crlr-Cre mice, it has been suggested that ventral CA3 pyramidal cells can be labeled by virus (Bernstein et al., 2020; Jinde et al., 2012). Therefore, the potential expression of virus in CA3 pyramidal cells was important to consider. It was particularly important because area CA3 pyramidal cells project to the DG, although the axon terminals are mainly the hilus (Ishizuka et al., 1990; Scharfman, 2007a; Scharfman and Myers, 2012). Nevertheless, it has been reported that temporal CA3 pyramidal cells innervate the GCs by axons in the DG IML (Li et al., 1994). Therefore, we examined the possibility that some of the IML axons we visualized in the IML or even the MML represented the axons of CA3 pyramidal cells, rather than MCs. We saw no evidence that CA3 axon terminals were localized to the IML or MML.

There are several implications of these findings. For example, the dorsal MCs have a much broader area of the dendrites of ventral GCs that they innervate compared to any GCs they target dorsally. Also, ventral MCs almost exclusively innervate the IML. GABAergic neurons that MCs innervate would have a similarly broad area for potential MC synapses from dorsal GCs but a more restricted area for MC synapses made by ventral MCs. There also is more potential for axon-axon, glial, or other interactions in the ML for dorsal MCs than ventral MCs.

If one only considers GCs, one would expect a greater potential for dorsal MCs to influence ventral and contralateral GCs by contacting more of the dendritic tree, and more opportunity to influence afferents to the GCs that lie in the MML. A functional interaction with the perforant path seems like an interesting possibility, although recent electrophysiological data suggest little direct interaction (Bernstein et al., 2020).

The reason that differences in dorsal and ventral MCs are important is based on the past reports that the DG exhibits significant functional differences in dorsal and ventral regions. Some of these studies suggest that the dorsal DG has functions related to cognition and spatial navigation, whereas ventral DG has functions related to contextual conditioning, mood, and anxiety (Kheirbek and Hen, 2011). If dorsal MCs have a broader IML plexus ventrally than dorsally, they may have significantly different effects on the GCs they target dorsally vs. ventrally. This could give them a greater range of effects near and far from their cell bodies. In contrast, ventral MCs may have very similar effects on the GCs they target, regardless of the position of the targeted cells in dorsal or ventral DG. As a result, the different projections across the septotemporal axis could give dorsal MCs the additional ability to encode information with a variable septotemporal valence. On the other hand, ventral MCs may have a more consistent, homogeneous function.

### 4.1.3 MC axons are heterotopic rather than homotopic in the contralateral DG

Studies from the 1980’s and 1990’s based on markers such as phaseolus vulgaris leucoagglutinin (PHAL), mainly in the rat, suggested that the axons of MCs were mainly destined for the IML in the ipsilateral hippocampus, terminating distal to the MC body (Scharfman and Myers, 2012). In addition, there was a homotopic distribution contralaterally, so dorsal MCs would project to the contralateral dorsal IML and ventral MCs would project to the contralateral ventral IML (Scharfman and Myers, 2012).

Since that time, no evidence has been provided that contradicts this idea of a homotopic contralateral projection. As a result, it is significant that the data in the present study show that MCs not only project homotopically in the contralateral DG, but also to heterotopic locations. Thus, a dorsal MC will project to distal ipsilateral locations, and to the majority of the septotemporal axis contralaterally. The exception could be the most ventral pole of the contralateral DG, because we found labeling relatively sparse in those locations.

Similarly, a ventral MC will project to the majority of the contralateral DG. Here the dorsal and ventral MCs may differ slightly because we found dorsal MCs projected to less of the septotemporal axis of the contralateral DG than ventral MCs. Together the data from dorsal and ventral MCs suggests a heterotopic distribution of the MC axon contralateral to its cell body and additional evidence that the dorsal and ventral MCs have a different axonal projection.

Why the present study found evidence of extensive contralateral projections compared to past studies is likely to be due to technical reasons. Thus, the viral expression of an opsin is membrane bound, whereas extracellular markers like PHAL, or intracellular markers such as biocytin are primarily cytoplasmic. With the ability to label the plasma membrane, virally-expressed opsins are able to make distal parts of axons and dendrites easier to visualize because the cytoplasm of these fine processes is small relative to the membrane.

The significance of the more widespread contralateral projection is interesting to consider. One possibility is that a more widespread axon makes MCs able to interconnect more lamellae of the DG. As such, MCs are more likely to serve roles that have been suggested for them before, such as a role as a sentinel cell, “broadcasting” its input to numerous GCs at almost all levels of the DG (Scharfman, 2016). The idea that MCs detect what is novel about the environment and send that to GCs so that environmental context can be processed has been suggested (Bernstein et al., 2019; Duffy et al., 2013), and could make it important for MCs to send their axons to all parts of the DG.

### 4.1.4 Blade differences

The distinctions between the dorsal and ventral MC axons were evident when measurements were made for the terminal fields in the dorsal blade, crest, and ventral blade (also referred to as the suprapyramidal blade, apex, and infrapyramidal blade respectively). The importance of these differences are not clear, although more and more is being detected that is distinct about the dorsal and ventral blades (Chawla et al., 2005; Scharfman et al., 2002; Schmidt et al., 2012).

## 5. LIMITATIONS

Diverse mouse strains, diverse ages and many endocrinological groups were not tested. As such, different ages and mouse strains could differ from the results shown here. Also, sex differences may exist if more detailed endocrinological studies of males and females were made. Sex differences are notable because of prior publications about sex differences in MCs (Guidi et al., 2006) and because a recent study showed that sex differences do appear to exist in the effects of MCs (Botterill et al., 2020).

## 6. CONCLUSIONS

The results show differences in dorsal and ventral MCs of the adult C57Bl6 mouse that is due to a broader terminal plexus in the distal axon projections of dorsal but not ventral MCs. The findings were thoroughly tested to confirm their reproducibility and lack of confounding factors. The implications are that the dorsal MCs may influence processing of information in the DG differently than ventral MCs. Dorsal-ventral differences in MCs could therefore contribute to dorsal-ventral differences of the DG.

## 7. DATA AVAILABILITY STATEMENT

Furthermore information and requests for reagents or resources should be directed to and will be fulfilled by the corresponding author, Dr. Helen Scharfman.

## 8. CONFLICTS OF INTREST

The authors declare that the research was conducted in absence of any commercial or financial interests that could be construed as a potential conflict of interest

## 9. AUTHOR CONTRIBUTIONS

*Conceptualization:* JJB, HES. *Data collection and analysis:* JJB, KJG, KYV, DAG. *Wrote the manuscript:* JJB & HES. All authors reviewed and approved the manuscript.

---

**Figure S1.**
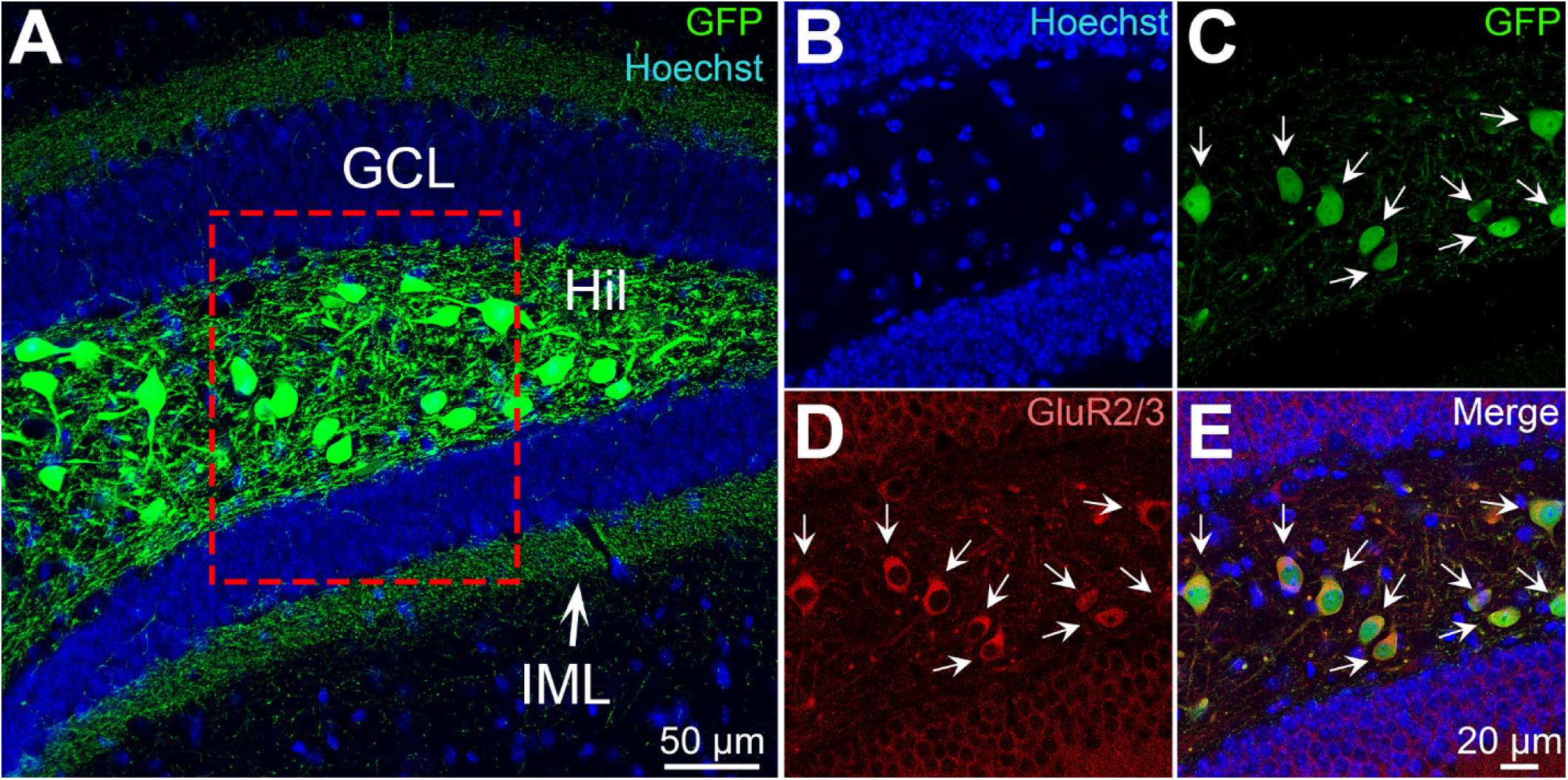
GFP cells colocalize with GluR2/3. **(A)** Representative photomicrograph showing GFP immunofluorescence in the hilus and IML. Insets show **(B)** Hoechst, **(C)** GFP and **(D)** GluR2/3. **(E)** Merged image shows that the GFP hilar cells strongly colocalize with GluR2/3 immunofluorescence, consistent with previous reports by our laboratory (Bernstein et al., 2020; Botterill et al., 2019) and others (Danielson et al., 2017; Jung et al., 2019; Yeh et al., 2018). (IML) inner molecular layer, (HIL) hilus, (GCL) granule cell layer.

**Figure S2.**
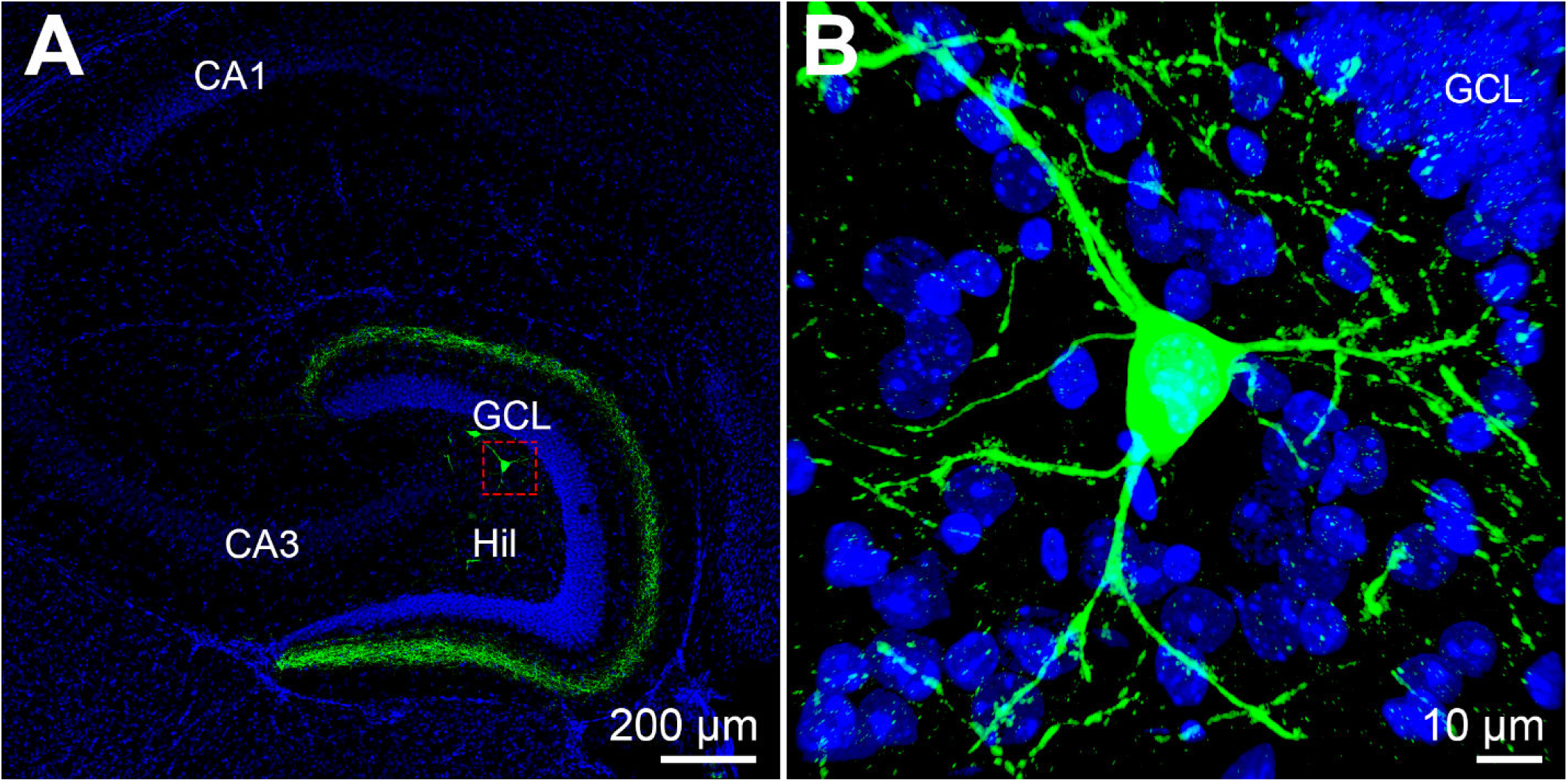
GFP cells in the non-injected hemisphere. **(A**) Photomicrograph from Figure 2C4 representing commissural projections in the non-injected hemisphere following a single dorsal hilus injection. GFP axons are observed distal to the GCL border in the ML. Several GFP cell bodies are also labeled. **(B)** High resolution z-stack of the GFP hilar cell in outlined in Panel A has morphological features consistent with a MC. (HIL) hilus, (GCL) granule cell layer.

**Figure S3.**
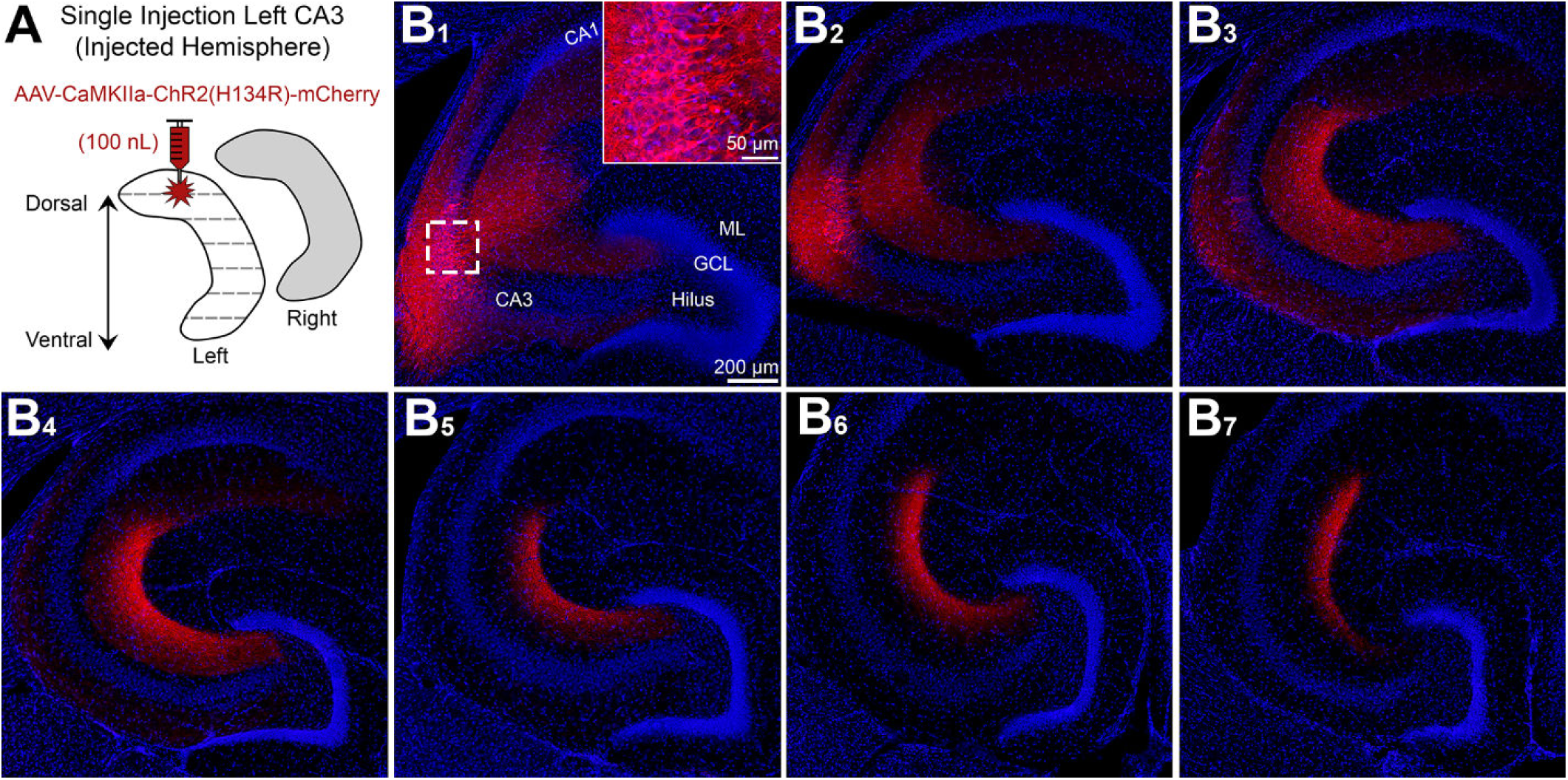
Viral expression in CA3 does not cause ML expression. (A) Viral injection schematic. AAV-CAMKIIa-ChR2(H134R)-mCherry was injected into the left dorsal CA3. **(B1-B2)** Proximal to the injection site, CA3 pyramidal cells in CA3A/B were labeled (see inset). (B3-B7). Distal to the injection site there were no mCherry expressing cells in the CA3 pyramidal layer, however a band of mCherry axons were observed in the CA3 stratum radiatum. Importantly, CA3 pyramidal cells caused no mCherry immunofluorescence in the molecular layer, which suggests that the ML immunofluorescence in MC targeted experiments was not due to CA3 contamination. (ML) molecular layer, (HIL) hilus, (GCL) granule cell layer.

**Figure S4.**
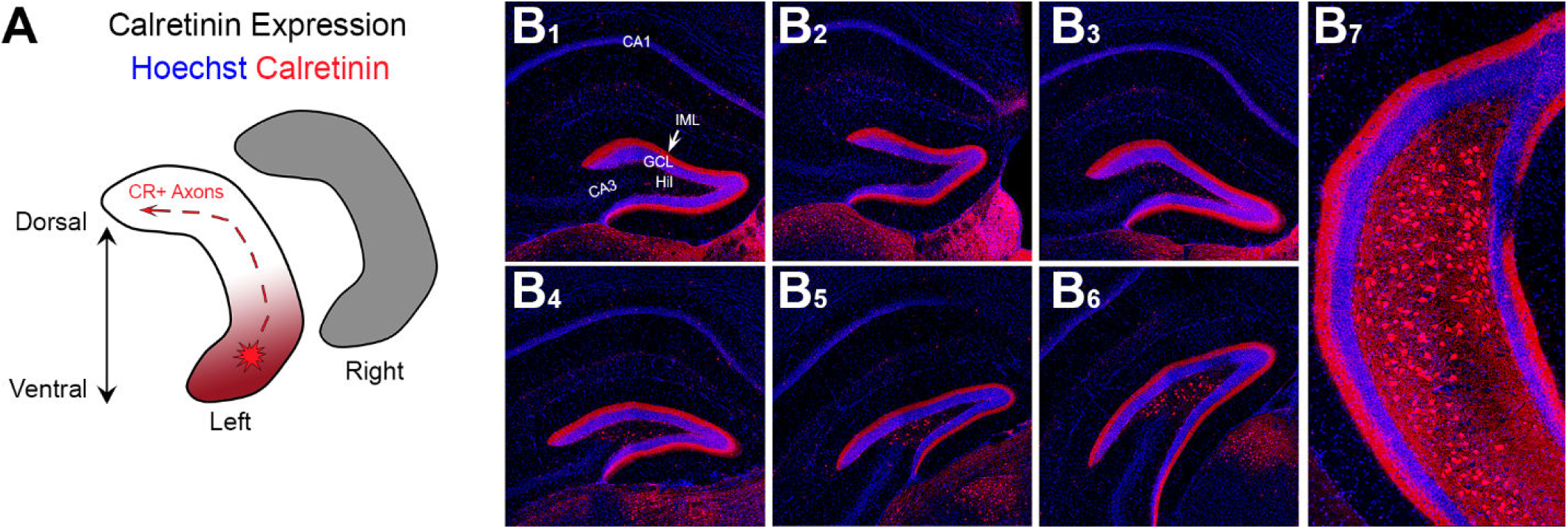
Calretinin immunoreactivity across the septotemporal axis of the DG. **(A)** Schematic of calretinin immunoreactivity. Within the DG, calretinin is primarily observed in ventral but not dorsal hilar cells. Calretinin also stains the IML throughout the entire septotemporal axis, presumably from calretinin-expressing MCs in the ventral DG. **(B1-B7)** Representative calretinin expression in the mouse DG. In the dorsal DG, there are few hilar cells and strong IML immunofluorescence. As sections proceed to more caudal regions and include the ventral DG, calretinin immunofluorescent hilar cells are observed as well as strong IML expression of calretinin. (IML) inner molecular layer, (HIL) hilus, (GCL) granule cell layer.

**Figure S5.**
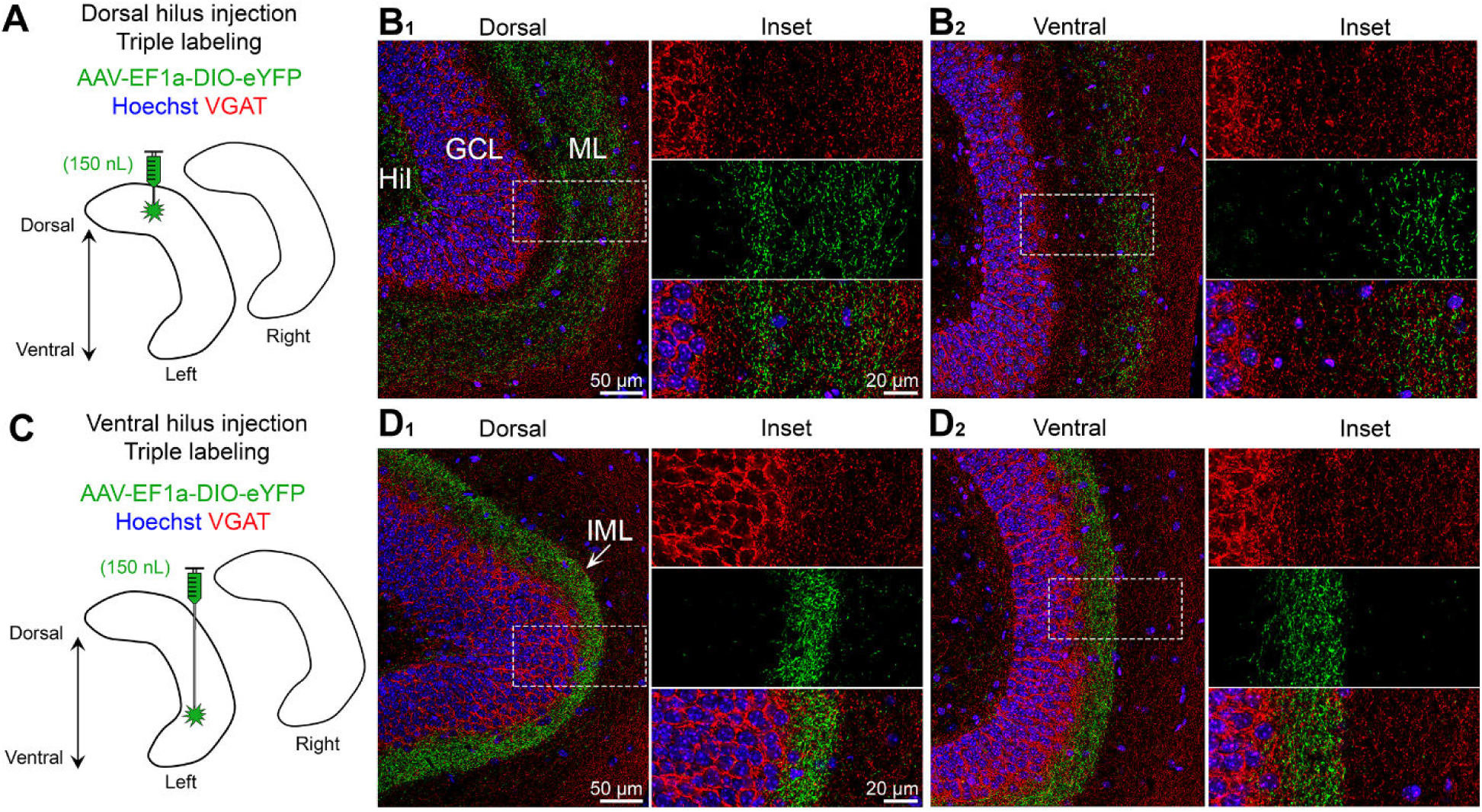
Dorsal and ventral MC axons show minimal colocalization with VGAT. **(A)** The left dorsal hilus was injected with AAV-EF1a-DIO-eYFP. **(B1-B2)** In dorsal and ventral sections, the GFP axon failed to show colocalized VGAT in any part of the molecular layer (ML). **(C)** The left ventral hilus was injected with AAV-EF1a-DIO-eYFP. **(D1-D2)** In dorsal and ventral horizontal sections, the GFP axon was restricted to the IML and showed minimal colocalization with VGAT. (I) inner (ML) molecular layer, (HIL) hilus, (GCL) granule cell layer.

**Figure S6.**
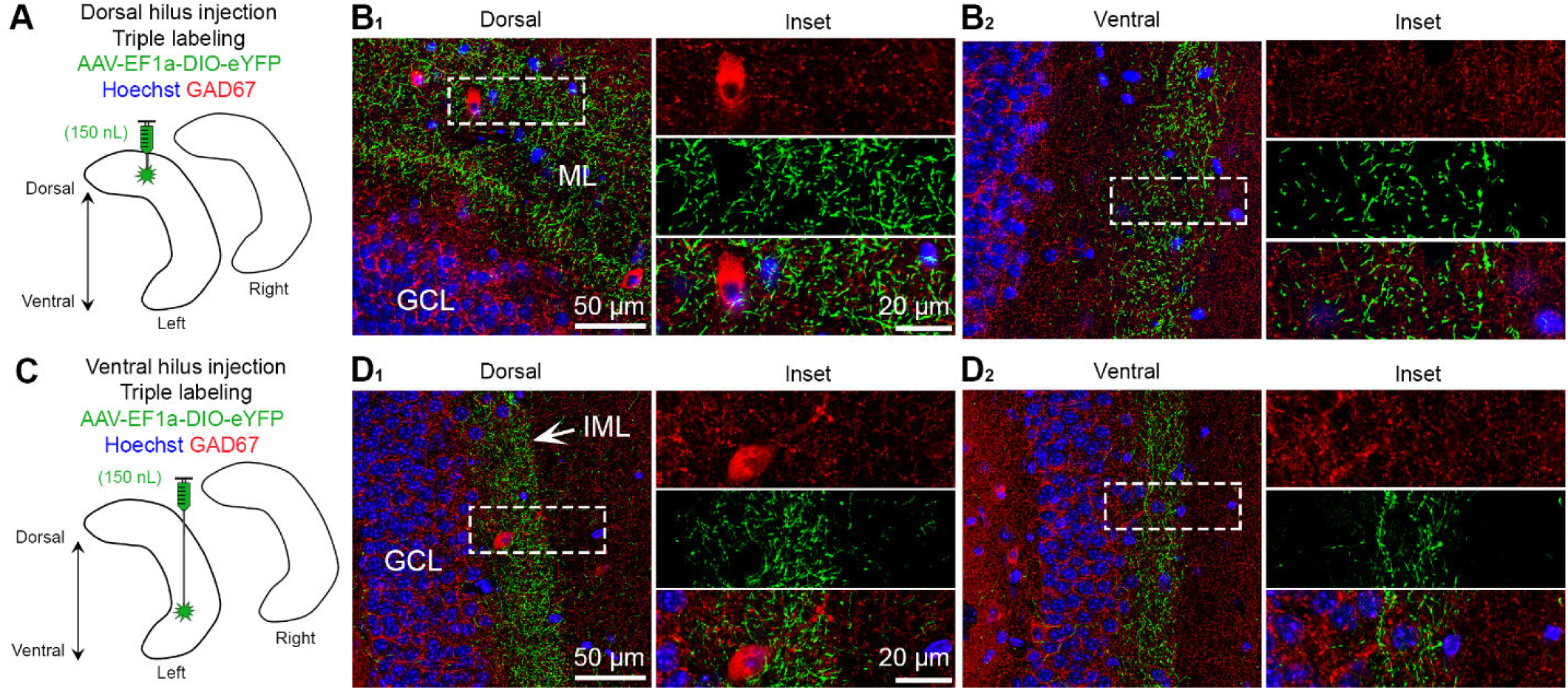
Dorsal and ventral MC axons show minimal colocalization with GAD67. **(A)** The left dorsal hilus was injected with AAV-EF1a-DIO-eYFP. **(B1-B2)** In dorsal and ventral sections, the GFP axon showed no detectable colocalization with GAD67 in any part of the molecular layer (ML). **(C)** The left ventral hilus was injected with AAV-EF1a-DIO-eYFP. **(D1-D2)** In dorsal and ventral horizontal sections, the GFP axon was restricted to the IML and showed minimal colocalization with GAD67. (I) inner (ML) molecular layer, (HIL) hilus, (GCL) granule cell layer.

**Supplemental Figure 7.**
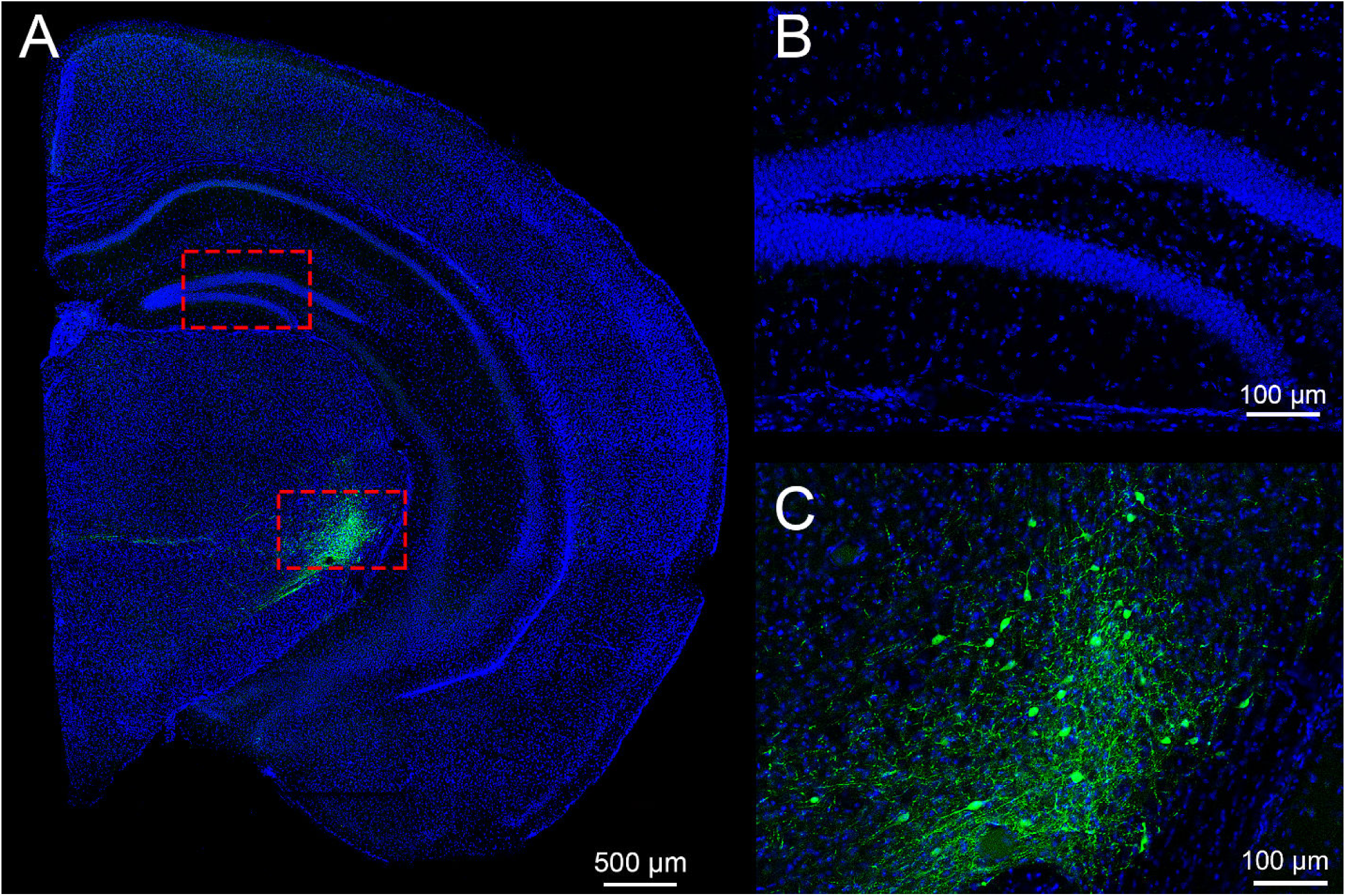
Incorrect targeting does not cause viral expression in the DG. **(A)** A mistargeted injection shows GFP expression in the thalamus. **(B)** Inset of the DG from Panel A shows there was no GFP expression in the hilus or IML. **(C)** High resolution inset of the GFP signal highlighted in Panel A. Numerous GFP cells were labeled.

## References

Amaral, D.G., Scharfman, H.E., and Lavenex, P. (2007). The dentate gyrus: fundamental neuroanatomical organization (dentate gyrus for dummies). Prog Brain Res 163, 3–22.

Armstrong, C., Szabadics, J., Tamas, G., and Soltesz, I. (2011). Neurogliaform cells in the molecular layer of the dentate gyrus as feed-forward gamma-aminobutyric acidergic modulators of entorhinal-hippocampal interplay. J Comp Neurol 519, 1476–1491.

Azevedo, E.P., Pomeranz, L., Cheng, J., Schneeberger, M., Vaughan, R., Stern, S.A., Tan, B., Doerig, K., Greengard, P., and Friedman, J.M. (2019). A role of Drd2 hippocampal neurons in context-dependent food intake. Neuron 102, 873–886 e875.

Bernstein, H.L., Lu, Y.L., Botterill, J.J., Duffy, A.M., LaFrancois, J., and Scharfman, H. (2020). Excitatory effects of dentate gyrus mossy cells and their ability to influence granule cell firing: an optogenetic study in adult mouse hippocampal slices. bioRxiv https://doiorg/101101/20200606137844.

Bernstein, H.L., Lu, Y.L., Botterill, J.J., and Scharfman, H.E. (2019). Novelty and novel objects increase c-Fos immunoreactivity in mossy cells in the mouse dentate gyrus. Neural Plast 2019, 1815371.

Blasco-Ibanez, J.M., and Freund, T.F. (1997). Distribution, ultrastructure, and connectivity of calretinin-immunoreactive mossy cells of the mouse dentate gyrus. Hippocampus 7, 307–320.

Botterill, J.J., Guskjolen, A.J., Marks, W.N., Caruncho, H.J., and Kalynchuk, L.E. (2015). Limbic but not non-limbic kindling impairs conditioned fear and promotes plasticity of NPY and its Y2 receptor. Brain Struct Funct 220, 3641–3655.

Botterill, J.J., Lu, Y.L., LaFrancois, J.J., Bernstein, H.L., Alcantara-Gonzalez, D., Jain, S., Leary, P., and Scharfman, H.E. (2019). An excitatory and epileptogenic effect of dentate gyrus mossy cells in a mouse model of epilepsy. Cell Rep 29, 2875–2889 e2876.

Botterill, J.J., Nogovitsyn, N., Caruncho, H.J., and Kalynchuk, L.E. (2017). Selective plasticity of hippocampal GABAergic interneuron populations following kindling of different brain regions. J Comp Neurol 525, 389–406.

Botterill, J.J., Vinod, K.Y., Gerencer, K.J., Teixeira, C.M., LaFrancois, J., and Scharfman, H.E. (2020). Bidirectional regulation of cognitive and anxiety-like behaviors by dentate gyrus mossy cells in male and female mice. BioRxiv https://doiorg/101101/20200705188664.

Buckmaster, P.S., and Schwartzkroin, P.A. (1994). Hippocampal mossy cell function: a speculative view. Hippocampus 4, 393–402.

Buckmaster, P.S., Wenzel, H.J., Kunkel, D.D., and Schwartzkroin, P.A. (1996). Axon arbors and synaptic connections of hippocampal mossy cells in the rat in vivo. J Comp Neurol 366, 271–292.

Chawla, M.K., Guzowski, J.F., Ramirez-Amaya, V., Lipa, P., Hoffman, K.L., Marriott, L.K., Worley, P.F., McNaughton, B.L., and Barnes, C.A. (2005). Sparse, environmentally selective expression of Arc RNA in the upper blade of the rodent fascia dentata by brief spatial experience. Hippocampus 15, 579–586.

Chawla, M.K., Sutherland, V.L., Olson, K., McNaughton, B.L., and Barnes, C.A. (2018). Behavior-driven arc expression is reduced in all ventral hippocampal subfields compared to CA1, CA3, and dentate gyrus in rat dorsal hippocampus. Hippocampus 28, 178–185.

Danielson, N.B., Turi, G.F., Ladow, M., Chavlis, S., Petrantonakis, P.C., Poirazi, P., and Losonczy, A. (2017). In vivo imaging of dentate gyrus mossy cells in behaving mice. Neuron 93, 552–559 e554.

Deller, T., and Leranth, C. (1990). Synaptic connections of neuropeptide Y (NPY) immunoreactive neurons in the hilar area of the rat hippocampus. J Comp Neurol 300, 433–447.

Duffy, A.M., Schaner, M.J., Chin, J., and Scharfman, H.E. (2013). Expression of c-fos in hilar mossy cells of the dentate gyrus in vivo. Hippocampus 23, 649–655.

Eyre, M.D., and Bartos, M. (2019). Somatostatin-expressing interneurons form axonal projections to the contralateral hippocampus. Front Neural Circuits 13, 56.

Fredes, F., Silva, M., Koppensteiner, P., Kobayashi, K., Joesch, M., and Shigemoto, R. (2019). Novelty gates memory formation through ventro-dorsal hippocampal interaction. BioRxiv https://doiorg/101101/673871.

Freund, T.F., and Buzsaki, G. (1996). Interneurons of the hippocampus. Hippocampus 6, 347–470.

Fujise, N., Liu, Y., Hori, N., and Kosaka, T. (1998). Distribution of calretinin immunoreactivity in the mouse dentate gyrus: II. Mossy cells, with special reference to their dorsoventral difference in calretinin immunoreactivity. Neuroscience 82, 181–200.

Guidi, S., Severi, S., Ciani, E., and Bartesaghi, R. (2006). Sex differences in the hilar mossy cells of the guinea-pig before puberty. Neuroscience 139, 565–576.

Haery, L., Deverman, B.E., Matho, K.S., Cetin, A., Woodard, K., Cepko, C., Guerin, K.I., Rego, M.A., Ersing, I., Bachle, S.M., et al. (2019). Adeno-associated virus technologies and Mmthods for targeted neuronal manipulation. Front Neuroanat 13, 93.

Halasy, K., and Somogyi, P. (1993). Subdivisions in the multiple GABAergic innervation of granule cells in the dentate gyrus of the rat hippocampus. Eur J Neurosci 5, 411–429.

Han, Z.S., Buhl, E.H., Lorinczi, Z., and Somogyi, P. (1993). A high degree of spatial selectivity in the axonal and dendritic domains of physiologically identified local-circuit neurons in the dentate gyrus of the rat hippocampus. Eur J Neurosci 5, 395–410.

Houser, C.R. (2007). Interneurons of the dentate gyrus: an overview of cell types, terminal fields and neurochemical identity. Prog Brain Res 163, 217–232.

Ishizuka, N., Weber, J., and Amaral, D.G. (1990). Organization of intrahippocampal projections originating from CA3 pyramidal cells in the rat. J Comp Neurol 295, 580–623.

Jinde, S., Zsiros, V., Jiang, Z., Nakao, K., Pickel, J., Kohno, K., Belforte, J.E., and Nakazawa, K. (2012). Hilar mossy cell degeneration causes transient dentate granule cell hyperexcitability and impaired pattern separation. Neuron 76, 1189–1200.

Jinno, S., Ishizuka, S., and Kosaka, T. (2003). Ionic currents underlying rhythmic bursting of ventral mossy cells in the developing mouse dentate gyrus. Eur J Neurosci 17, 1338–1354.

Jung, D., Kim, S., Sariev, A., Sharif, F., Kim, D., and Royer, S. (2019). Dentate granule and mossy cells exhibit distinct spatiotemporal responses to local change in a one-dimensional landscape of visual-tactile cues. Sci Rep 9, 9545.

Kheirbek, M.A., Drew, L.J., Burghardt, N.S., Costantini, D.O., Tannenholz, L., Ahmari, S.E., Zeng, H., Fenton, A.A., and Hen, R. (2013). Differential control of learning and anxiety along the dorsoventral axis of the dentate gyrus. Neuron 77, 955–968.

Kheirbek, M.A., and Hen, R. (2011). Dorsal vs ventral hippocampal neurogenesis: implications for cognition and mood. Neuropsychopharmacology 36, 373–374.

Kosaka, T., Katsumaru, H., Hama, K., Wu, J.Y., and Heizmann, C.W. (1987). GABAergic neurons containing the Ca2+-binding protein parvalbumin in the rat hippocampus and dentate gyrus. Brain Res 419, 119–130.

Lanciego, J.L., and Wouterlood, F.G. (2020). Neuroanatomical tract-tracing techniques that did go viral. Brain Struct Funct 225, 1193–1224.

Li, X.G., Somogyi, P., Ylinen, A., and Buzsaki, G. (1994). The hippocampal CA3 network: an in vivo intracellular labeling study. J Comp Neurol 339, 181–208.

Myers, C.E., and Scharfman, H.E. (2009). A role for hilar cells in pattern separation in the dentate gyrus: a computational approach. Hippocampus 19, 321–337.

Oh, S.J., Cheng, J., Jang, J.H., Arace, J., Jeong, M., Shin, C.H., Park, J., Jin, J., Greengard, P., and Oh, Y.S. (2019). Hippocampal mossy cell involvement in behavioral and neurogenic responses to chronic antidepressant treatment. Mol Psychiatry.

Puighermanal, E., Biever, A., Espallergues, J., Gangarossa, G., De Bundel, D., and Valjent, E. (2015). drd2-cre:ribotag mouse line unravels the possible diversity of dopamine d2 receptor-expressing cells of the dorsal mouse hippocampus. Hippocampus 25, 858–875.

Scharfman, H.E. (2007a). The CA3 “backprojection” to the dentate gyrus. Prog Brain Res 163, 627–637.

Scharfman, H.E. (2007b). The dentate gyrus: a comprehensive guide to structure, function, and clinical implications, Vol 163 (Elsevier).

Scharfman, H.E. (2016). The enigmatic mossy cell of the dentate gyrus. Nat Rev Neurosci 17, 562–575.

Scharfman, H.E. (2017). Advances in understanding hilar mossy cells of the dentate gyrus. Cell Tissue Res.

Scharfman, H.E., and Myers, C.E. (2012). Hilar mossy cells of the dentate gyrus: a historical perspective. Front Neural Circuits 6, 106.

Scharfman, H.E., Sollas, A.L., Smith, K.L., Jackson, M.B., and Goodman, J.H. (2002). Structural and functional asymmetry in the normal and epileptic rat dentate gyrus. J Comp Neurol 454, 424–439.

Schmidt, B., Marrone, D.F., and Markus, E.J. (2012). Disambiguating the similar: the dentate gyrus and pattern separation. Behav Brain Res 226, 56–65.

Senzai, Y., and Buzsaki, G. (2017). Physiological properties and behavioral correlates of hippocampal granule cells and mossy cells. Neuron 93, 691–704.

Sperk, G., Hamilton, T., and Colmers, W.F. (2007). Neuropeptide Y in the dentate gyrus. Prog Brain Res 163, 285–297.

Yeh, C.Y., Asrican, B., Moss, J., Quintanilla, L.J., He, T., Mao, X., Casse, F., Gebara, E., Bao, H., Lu, W., et al. (2018). Mossy cells control adult neural stem cell quiescence and maintenance through a dynamic balance between direct and indirect pathways. Neuron 99, 493–510 e494.

